# Interferons and viruses induce a novel primate-specific isoform dACE2 and not the SARS-CoV-2 receptor ACE2

**DOI:** 10.1101/2020.07.19.210955

**Authors:** Olusegun O. Onabajo, A. Rouf Banday, Wusheng Yan, Adeola Obajemu, Megan L. Stanifer, Deanna M. Santer, Oscar Florez-Vargas, Helen Piontkivska, Joselin Vargas, Carmon Kee, D. Lorne J. Tyrrell, Juan L. Mendoza, Steeve Boulant, Ludmila Prokunina-Olsson

**Affiliations:** Laboratory of Translational Genomics, Division of Cancer Epidemiology and Genetics, National Cancer Institute, National Institutes of Health, Bethesda, MD, USA; Department of Infectious Diseases, Molecular Virology, University Hospital Heidelberg, Heidelberg, Germany; Li Ka Shing Institute of Virology and Department of Medical Microbiology and Immunology, University of Alberta, Edmonton, Alberta, Canada; Department of Biological Sciences and Brain Health Research Institute, Kent State University, Kent, OH, USA; Pritzker School of Molecular Engineering and Department of Biochemistry and Molecular Biology, University of Chicago, Chicago, IL, USA; Division of Cellular Polarity and Viral Infection, German Cancer Research Center (DKFZ), Heidelberg, Germany; Department of Infectious Diseases, Virology, University Hospital Heidelberg, Heidelberg, Germany

**Keywords:** ACE2, Interferon-stimulated gene, Interferon, SARS-CoV-2, COVID-19, immune response

## Abstract

Severe acute respiratory syndrome coronavirus-2 (SARS-CoV-2), which causes COVID-19, utilizes angiotensin-converting enzyme 2 (ACE2) for entry into target cells. *ACE2* has been proposed as an interferon-stimulated gene (ISG). Thus, interferon-induced variability in *ACE2* expression levels could be important for susceptibility to COVID-19 or its outcomes. Here, we report the discovery of a novel, primate-specific isoform of *ACE2*, which we designate as *deltaACE2 (dACE2)*. We demonstrate that *dACE2*, but not *ACE2*, is an ISG. *In vitro*, dACE2, which lacks 356 N-terminal amino acids, was non-functional in binding the SARS-CoV-2 spike protein and as a carboxypeptidase. Our results reconcile current knowledge on *ACE2* expression and suggest that the ISG-type induction of *dACE2* in IFN-high conditions created by treatments, inflammatory tumor microenvironment, or viral co-infections is unlikely to affect the cellular entry of SARS-CoV-2 and promote infection.

## INTRODUCTION

Cells expressing *ACE2* are potential targets of SARS-CoV-2 infection^1,2^. Studies based on single-cell RNA-sequencing (scRNA-seq) of lung cells have identified type II pneumocytes, ciliated cells, and transient secretory cells as the main types of *ACE2-*expressing cells^3,4^. Furthermore, *ACE2* was proposed to be an ISG, based on its inducible expression in cells treated with interferons (IFNs) or infected by viruses that induce IFN responses, such as influenza^4,5^. These findings implied that the induction of *ACE2* expression in IFN-high conditions could result in an amplified risk of SARS-CoV-2 infection^4,5^. Concerns could also be raised about possible *ACE2*-inducing side effects of IFN-based treatments proposed for COVID-19^6–9^.

ACE2 plays multiple roles in normal physiological conditions and as part of host tissue-protective machinery during damaging conditions, including viral infections. As a terminal carboxypeptidase, ACE2 cleaves a single C-terminal residue from peptide hormones such as angiotensin II and des-Arg9-bradykinin. Angiotensin II is generated from its precursor angiotensin I by ACE, and is proteolytically converted into another biologically active peptide, angiotensin-(1-7) by ACE2^10,11^. ACE and ACE2 belong to the renin-angiotensin-aldosterone system, which regulates blood pressure and fluid-electrolyte balance; dysfunction of this system contributes to comorbidities in COVID-19^10,12^. des-Arg9-bradykinin is generated from bradykinin and belongs to the kallikrein-kinin system, which is critical in regulating vascular leakage and pulmonary edema, early signs of severe COVID-19^13,14^.

High plasma angiotensin II levels were found responsible for coronavirus-associated acute respiratory distress syndrome (ARDS), lung damage and high mortality in mouse models^15,16^ and as a predictor of lethality in avian influenza in humans^17,18^. ACE2, which decreases the levels of angiotensin II, was identified as a protective factor in the same conditions. The hijacking of the normal host tissue-protective machinery guarded by ACE2 was suggested as a mechanism through which SARS-CoV could infect more cells^4,5^. Thus, it is critically important to identify factors affecting *ACE2* expression in normal physiological processes and during viral infections and associated pathologies, such as in COVID-19.

Herein, aiming to explore the IFN-induced expression of *ACE2* and its role in SARS-CoV-2 infection, we identified a novel, primate-specific isoform of *ACE2*, which we designate as *deltaACE2* (*dACE2*). We then showed that *dACE2,* but not *ACE2,* is induced in various human cell types by IFNs and viruses; this information is important to consider for future therapeutic strategies and understanding susceptibility and outcomes of COVID-19.

## RESULTS

### *dACE2* is a novel inducible and primate-specific isoform of *ACE2*

To address the extent to which IFNs induce the expression of *ACE2* in human cells, we used our existing RNA-seq dataset (NCBI SRA: PRJNA512015) of a breast cancer cell line T47D infected with Sendai virus (SeV), known to be a strong inducer of IFNs and ISGs^19–21^. Accordingly, no IFNs were expressed in T47D cells at baseline, but SeV strongly induced *IFNB1,* a type I IFN and all type III IFNs (*IFNL1, 2, 3,* and *4*). Several well-known ISGs (*ISG15*, *MX1* and *IFIT1*) were moderately expressed at baseline but were strongly induced by SeV (**Table S1**). *ACE2* was not expressed at baseline but was strongly induced by SeV, exclusively as an isoform initiated from a novel first exon in intron 9 of the full-length *ACE2* gene (**Figure 1A, B**).

**Figure 1.**
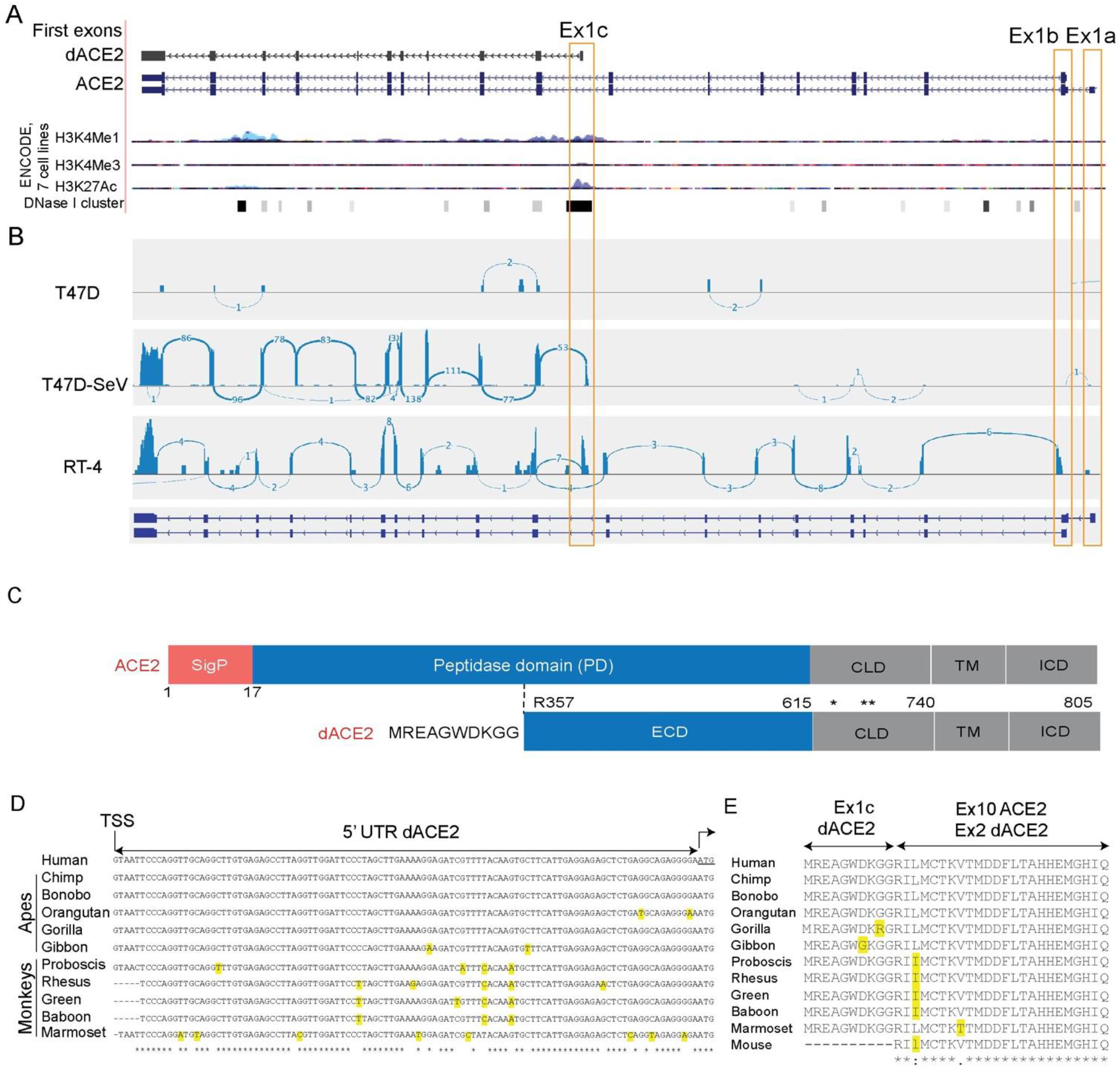
*dACE2* is a novel primate-specific and virally-induced isoform of *ACE2.* **A)** UCSC genome browser view of the human *ACE2* region (chrX:15,560,521-15,602,580, GRCh38/hg38) showing alternative first exons *ACE2*-Ex1a and Ex1b and a novel Ex1c that creates a short isoform designated as delta *ACE2* (*dACE2*). ENCODE epigenetic markers (H3K4me1, H3K4me3, H3K27ac, and cluster of DNase I hypersensitivity sites) indicate that *dACE2*-Ex1c is located within a putative regulatory region that can affect gene expression. **B)** RNA-seq Sashimi plots depict splicing patterns defining *ACE2* and *dACE2* isoforms in SeV/mock-infected T47D cells and uninfected RT-4 cells. **C)** ACE2 is a single-span transmembrane protein with a signal peptide (SigP) of 17 aa and four functional domains – peptidase domain (PD, aa 18-615), collectrin-like domain (CLD, aa 616-740), transmembrane domain (TM, aa 741-761), and intracellular domain (ICD, aa 762-805). In dACE2, no signal peptide is predicted; the extracellular domain (ECD) starts from aa R357; the first 356 aa are replaced by 10 aa of a unique protein sequence; * and ** - cleavage sites by membrane-bound proteases ADAM17 and TMPRSS2, respectively. **D)** Alignments of the *dACE2* 5’UTRs in primates. **E)** Alignments of protein sequences encoded by *dACE2*-Ex1c and part of the downstream exon. dACE2 is not predicted to be encoded in any non-primate species.

RNA-seq analysis in additional cell lines demonstrated that *ACE2* exists as two full-length transcripts initiated from two independent first exons, which we designated as Ex1a and Ex1b (the latter is shared between these transcripts). Additionally, a truncated transcript was initiated from this novel first exon in intron 9, which we designated as Ex1c (**Figure 1B**). The combination of ENCODE chromatin modification marks (H3K4me1, H3K4me3, H3K27ac, and a cluster of DNase I hypersensitivity sites, **Figure 1A**) suggests that Ex1c, but not Ex1a and Ex1b, is located within a putative regulatory region that might affect gene expression. Analysis of corresponding promoters for binding motifs for transcription factors relevant for IFN signaling predicted multiple motifs within the promoter of *ACE2-*Ex1c (P3) (**Figure S1**). In contrast, no ISG-type motifs were predicted in promoters of *ACE2*-Ex1a (P1) and Ex1b (P2), and only one ISG-type motif was predicted within the 5’UTR of Ex1b (**Figure S1**).

The novel *ACE2* isoform at 5p22.2 locus of human chromosome X is predicted to encode a protein of 459 aa, in which Ex1c encodes the first 10 aa, which are unique. Compared to the full-length ACE2 protein of 805 aa, the truncation eliminates 17 aa of the signal peptide (SigP) and 339 aa of the N-terminal peptidase domain (PD, **Figure 1C**). We designate this novel transcript as *deltaACE2* (*dACE2,* NCBI GenBank accession number MT505392). Of the 100 vertebrate species with genomic sequences available through the UCSC Genome Browser, the 140-bp sequence of *dACE2*-Ex1c is highly conserved only in primates (**Figure 1D, E** and **Figure S2A**, **B**).

### *dACE2* is induced by *in vitro* treatment with IFNs

We confirmed the SeV-induced expression of the full-length *dACE2* by RT-PCR (**Figure 2A, B**) and verified the corresponding PCR products by Sanger-sequencing. Using custom-designed assays, we explored *ACE2* and *dACE2* expression in multiple cell lines (**Figure 2C**, **Table S2A**). In most cell lines tested, *dACE2* but not *ACE2* was strongly upregulated by SeV infection (**Figure 2B, C**). To directly address whether IFN was responsible for the induced expression of *dACE2*, we performed expression analysis in primary normal human bronchial epithelial (NHBE) cells^22^ and human intestinal (colon and ileum) organoid cultures^23^. In NHBE cells from 5 healthy donors, baseline expression levels of *dACE2* and *ACE2* were comparable, but only *dACE2* was significantly induced by treatment with IFN-α or IFN-λ3 (**Figure 2D, Table S2B**). In contrast, *ACE2* was highly expressed already at baseline both in colon and ileum organoid cultures, while the expression of *dACE2* was very low. Treatments with IFN-β or a cocktail of IFNλ1-3 significantly induced only expression of *dACE2* and not *ACE2* (**Figure 2E, Table S2C**). In both cell models, the expression pattern of *dACE2* was similar to that of the known ISGs – *MX1* (**Figure 2D, Table S2B**) and *IFIT1* (**Figure 2E, Table S2C**).

**Figure 2.**
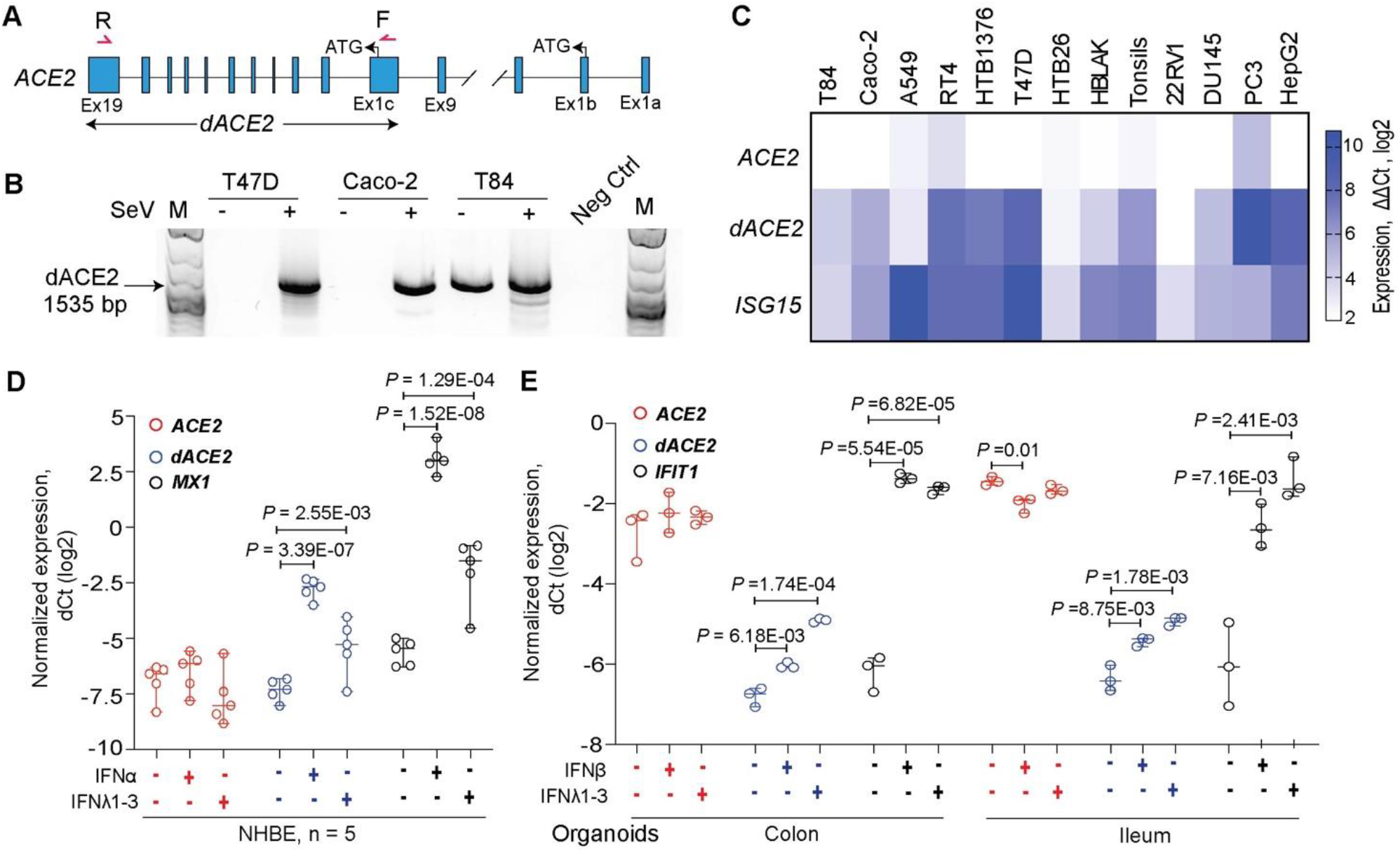
*dACE2* is induced by IFNs *in vitro.* **A)** Schematic representation of *ACE2* and *dACE2* transcripts and the position of the forward (F) and reverse (R) PCR primers to generate full-length *dACE2* amplicons. **B**) Agarose gel showing an RT-PCR product of 1535 bp corresponding to full-length *dACE2* in several cell lines with/without SeV infection. **C**) Heatmap of *ACE2* and *dACE2* expression and a positive control ISG (*ISG15*) measured by TaqMan expression assays in human cell lines infected with SeV for 12 hrs; colors represent expression differences as ddCt (log2) normalized by endogenous controls (*GAPDH* and *ACTB*) and comparing SeV-infected to uninfected samples. **D)** Expression of *ACE2* and *dACE2* and the positive control ISG *MX1* in primary normal human bronchial epithelial (NHBE) cells treated with IFNα or IFN-λ3 for 24 hrs vs. untreated cells. **E)** Expression of *ACE2* and *dACE2* and the positive control ISG *IFIT1* in colon and ileum organoid cultures from one donor; the organoids were treated with IFNβ or a cocktail of IFN-λ1-3 for 24 hrs. P-values are for Student’s T-test.

### *dACE2* is induced in virally infected human respiratory epithelial cells

To investigate whether *dACE2* expression is induced by RNA viruses, which are potent inducers of the IFN response, we *de novo* quantified the expression of *ACE2*-Ex1a, Ex1b, and the newly annotated *dACE2*-Ex1c in several public RNA-seq datasets of virally infected human respiratory epithelial cells. In an RNA-seq dataset of human nasal airway epithelial cells from asthmatic patients *ex-vivo* infected with respiratory rhinovirus strains RV-A16 and RV-C15 (NCBI SRA: PRJNA627860), both *ACE2* and *dACE2* were expressed (**Figure 3A**). Compared to uninfected cells, *dACE2*-Ex1c expression was strongly induced by both viruses – by RV-A16 (2.58-fold; P = 2.52E-13) and RV-C15 (2.42-fold; P = 2.52E-12), while expression of *ACE2*-Ex1b was weakly induced only by RV-C15 (1.13-fold; P = 0.032) (**Figure 3B**). Only *dACE2* expression strongly correlated with multiple ISGs and IFNs (**Figure 3C**). Similarly, in human lung explants infected with influenza A/H3N2 virus (NCBI SRA: PRJNA557257), only *dACE2* was induced by infection, and its expression correlated with the levels of IFNs and ISG (**Figure 3D, E**).

**Figure 3.**
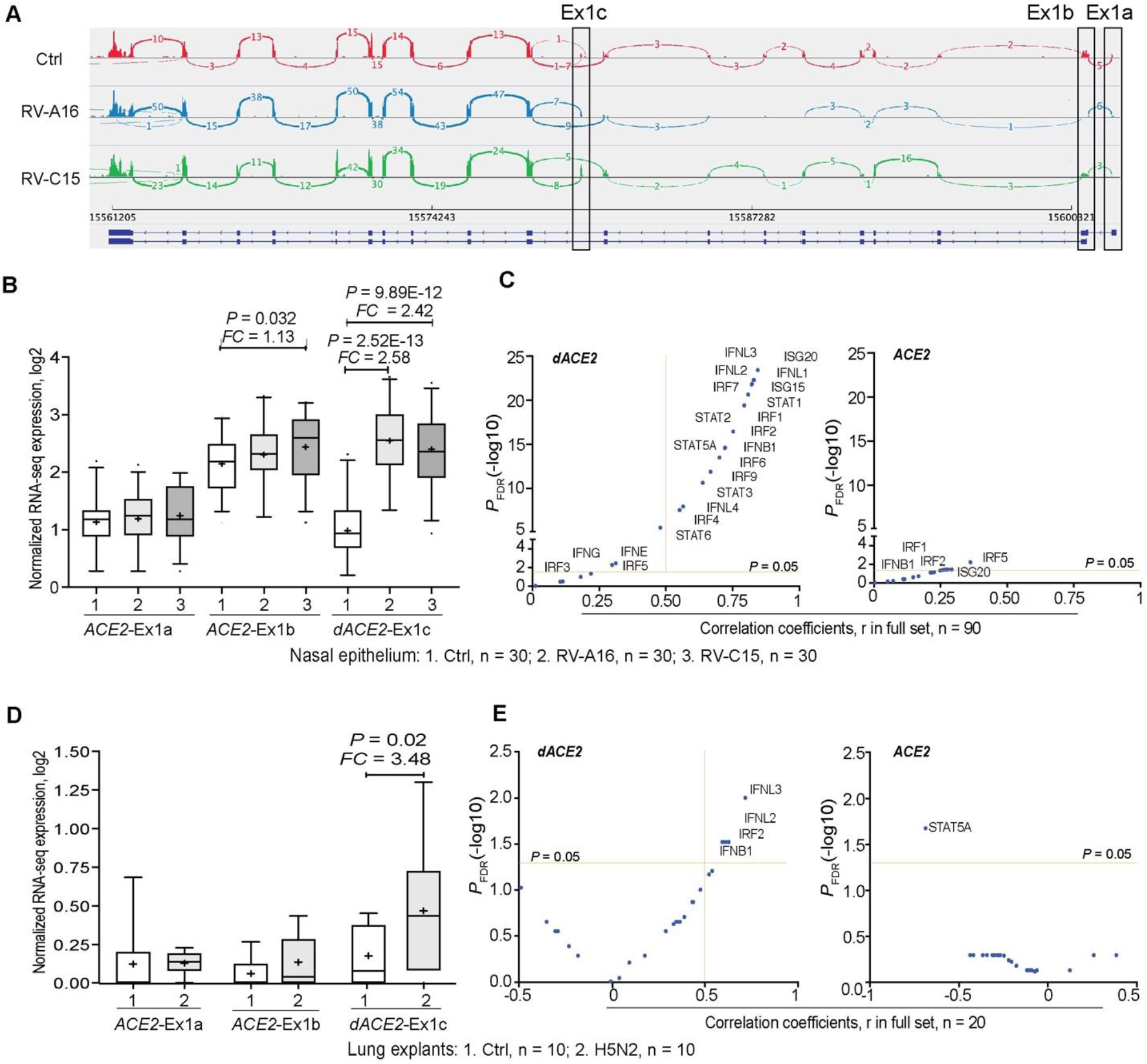
*dACE2* is induced in virally infected human respiratory cells. Expression patterns of *ACE2,* represented by Ex1a and Ex1b and *dACE2*, represented by Ex1c in **A-C)** mock (ctrl) and rhinovirus (RV)-infected human nasal epithelial cells and **D-E)** in uninfected (ctrl) and influenza H5N2-infected cells from human lung explants. **A)** RNA-seq Sashimi plots **B)** Expression levels **C)** Correlation coefficients with select ISGs and IFNs in the full dataset of 90 samples. **D)** Expression levels E) Correlation coefficients with select ISGs and IFNs in the full dataset of 20 samples. *FC* – fold change compared to mock. P-values are for the non-parametric Mann–Whitney U test.

It was reported that in contrast to *ACE2* expression in human cells, *Ace2* was not induced in primary mouse tracheal basal cells in response to *in vitro* and *in vivo* IFN-stimulation, and upon *in vivo* viral infection^4^. To explore this further, we analyzed a dataset for human and mouse lung cells infected with the respiratory syncytial virus (RSV, NCBI SRA: PRJNA588982). Indeed, we did not observe induction of *Ace2* in mouse lung cells (**Figure S3A**), while *dACE2* but not *ACE2* was induced in RSV-infected human pulmonary carcinoma cell line (H292) (**Figure S3B**). These results illustrate that the identification of *ACE2* as an ISG^4,5^ was likely based on the detection of inducible expression of *dACE2,* since 3’-scRNA-seq would detect both *ACE2* and *dACE2*. The significant differences between the human and mouse sequences corresponding to *dACE2*-Ex1c and corresponding promoters might be responsible for the lack of ISG-type *ACE2* expression in the mouse (**Figure 1D**, **Figure S2A, B**).

### *dACE2* is enriched in squamous epithelial tumors

We explored the expression patterns of *dACE2* in various human tissues. In a dataset of 95 normal human tissues of 27 types, *dACE2-*Ex1c was detectable in select tissues but at very low levels (≤ 10 RNA-seq reads), while *ACE2-*Ex1b expression was common (**Figure S4**). In the Genotype-Tissue Expression (GTEx) set of normal human tissues, only total gene expression was available for *ACE2,* with the highest expression observed in the testes and small intestine (**Figure S5**). We hypothesized that as an ISG, *dACE2* might be absent or expressed at low levels in normal tissues, but could be induced by the inflammatory tissue microenvironment. We explored the data from The Cancer Genome Atlas (TCGA), which represents the largest collection of tumors and tumor-adjacent normal tissues. Expression of both *ACE2* and *dACE2* was detectable in many tumor-adjacent normal tissues (**Figure 4A**, **Figure S6**). In the set of 10,185 TCGA tumors of 33 types, *ACE2*-Ex1a, Ex1b and *dACE2*-Ex1c were expressed in 12.6%, 38.0% and 16.8% of tumors, respectively, with ≥5 RNA-seq reads/sample. *dACE2* was most expressed in the sets of head and neck squamous carcinoma (HNSC) and lung squamous carcinoma (LUSC), which represent oral and bronchial mucosal epithelial surfaces, while *ACE2* was most expressed in kidney tumors (**Figure S7**).

**Figure 4.**
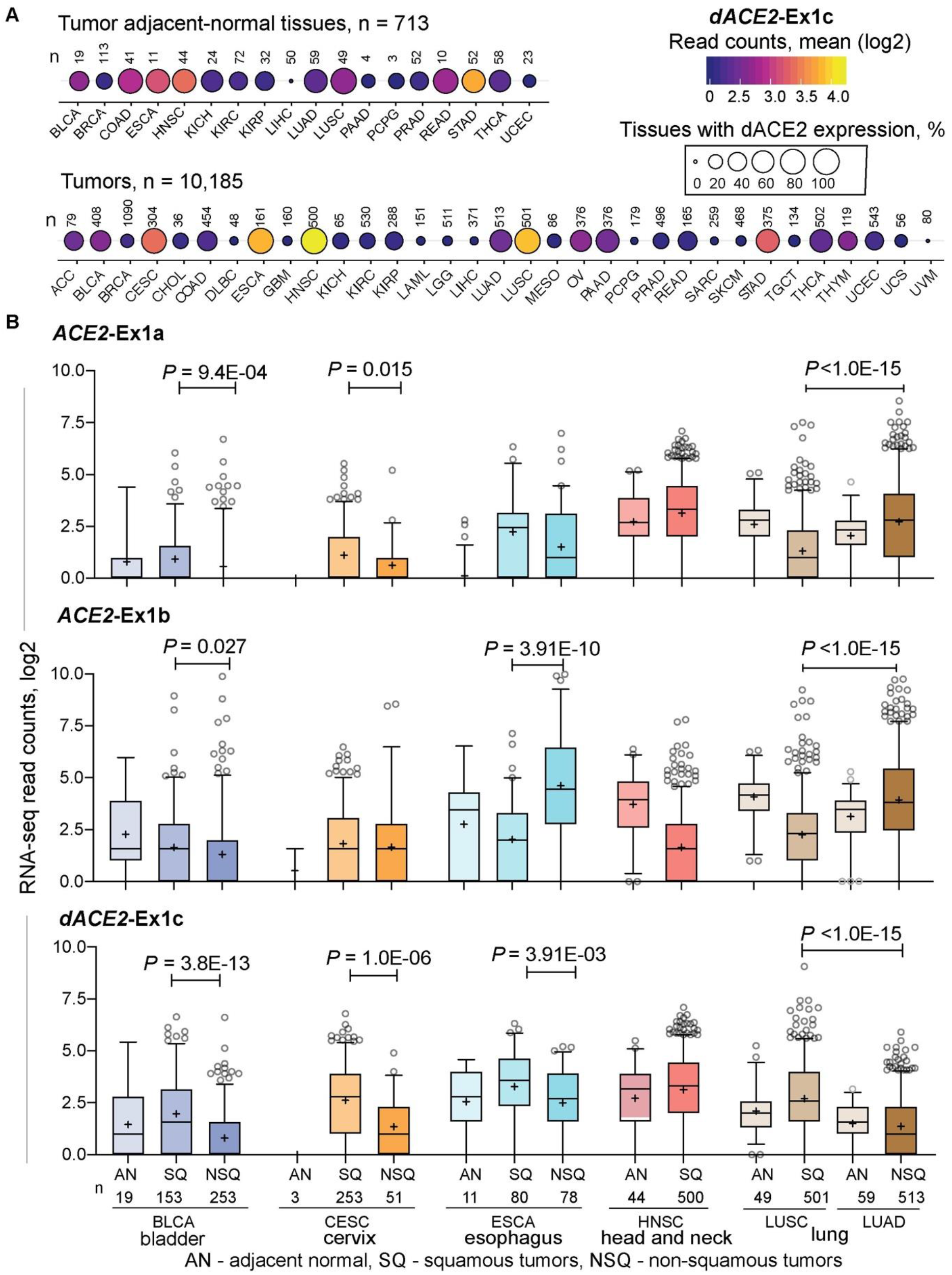
Expression of *dACE2* is enriched in squamous epithelial tumors. **A)**. Bubble plot showing mean expression levels (RNA-seq read counts) and proportions of samples (%) with *dACE2-*Ex1c in TCGA tumor-adjacent normal (AN) tissues and 33 tumor types. d *ACE2-*Ex1c is expressed at relatively high levels and in many tumors of the bladder (BLCA), cervix (CESC), esophagus (ESCA), head and neck (HNSC), and lung squamous carcinoma (LUSC). **B)** RNA-seq counts of *ACE2-*Ex1a and Ex1b and *dACE2-*Ex1c in tumor-adjacent normal (AN), squamous tumors (SQ) and non-squamous tumors (NSQ). Expression of *dACE2* is significantly higher in SQ compared to NSQ tumors of the same tissue origin and corresponding AN tissues. Specifically – *dACE2*-Ex1c is expressed similarly in AN tissues adjacent to LUSC and LUAD, while it is significantly higher in corresponding tumors and higher in LUSC than in LUAD, due to clonal origin of these tumors from cells with differential expression of *dACE2.* P-values are for the nonparametric Mann–Whitney U test.

Generally, *dACE2* expression was enriched in squamous tumors representing epithelial tracts. Squamous carcinomas of the lung (LUSC) and head and neck (HNSC) represent respiratory tract, esophageal cancer (ESCA) - upper gastrointestinal, and bladder cancer (BLCA) and cervical squamous carcinoma (CESC) - urogenital tract (**Figure 4A**). In each tumor type, *dACE2* expression was significantly higher in squamous compared to non-squamous tumors and adjacent-normal tissues (**Figure 4B**). Because tumors represent results of clonal expansion of individual cells, this analysis highlighted the differential etiology (squamous vs. non-squamous) of *ACE2-* and *dACE2* expressing cells in various tissues.

### *dACE2* expression is IFN-γ-inducible

In addition to cell type origin, the observed enrichment of *dACE2* expression in some tumors might reflect persistent IFN exposure due to an inflammatory tumor microenvironment or underlying infection. IFN-γ-induced signature emerged as a prominent feature in airway epithelial cells of COVID-19 patients, and this signature was linked with enrichment of cytotoxic T lymphocytes (CTLs)^5^. Unlike normal tissues, tumors are extensively infiltrated by immune cells, making TCGA dataset particularly informative for the analysis of IFN signatures. *IFNG* was the most commonly expressed IFN in TCGA tumors (with 61% of all tumors expressing *IFNG* at RSEM ≥ 1, mean RSEM = 19.8), while other IFNs were poorly expressed (mean RSEM ≤ 1.3, **Figure S8A**). d*ACE2-*Ex1c levels significantly correlated (P ≤ 0.01, r ≥ 0.2) with *IFNG* expression in 8 out of 32 tumor types tested (**Figure S8B**), while *ACE2-*Ex1b showed only moderate and predominantly negative correlations with *IFNG* expression in some tumor types (**Figure S8C**). Furthermore, *in vitro* treatment with IFN-γ significantly induced *dACE2,* but not *ACE2* (**Figure S8D**). Thus, in addition to type I and type III IFNs (**Figure 2E-G**), *dACE2* expression might be partly driven by IFN-γ contributed by tumor-infiltrating immune cells or inflamed virally infected tissues.

However, in TCGA-LUSC dataset (n=501), which represents *ACE2-* and *dACE2*-expressing tumors of bronchial origin (**Figure 4B**), *IFNG* expression did not correlate with *dACE2* (**Figure S8B**). Using a curated list of 270 ISGs^24^, we used an unsupervised machine learning approach to assign all LUSC tumors to 6 clusters based on their ISG expression profiles (**Figure S9A**, **B**). A set of ISGs (n=100) that most strongly contributed to the definition of these clusters was used for correlation analysis. The analysis in a cluster that included 114 LUSC tumors with the highest ISG expression showed that *dACE2* was strongly and significantly (FDR p-value < 0.05) correlated with expression of 20 ISGs and *ACE2 -* with 5 ISGs (**Figure S9C**, **D**). Thus, in LUSC tumors, *dACE2* was expressed as an ISG but unlikely driven by IFN-γ, suggesting multiple factors might be contributing to this expression.

### *dACE2* is induced by SARS-CoV-2 *in vitro*

Once we established that *dACE2* is an ISG in multiple human cells under various conditions, we tested whether its expression could also be induced by SARS-CoV-2. There was a noticeable difference in baseline expression levels of *ACE2* and *dACE2* in three cell lines tested (Calu3, Caco-2 and T84). Expression of *ACE2* and *dACE2* was much higher in a lung adenocarcinoma cell line Calu3 compared to both colon adenocarcinoma cell lines Caco2 and T84 (**Figure 5A, Table S2D**). Baseline *dACE2* expression in T84 was higher than in Caco-2, in line with the RT-PCR results (**Figure 1B)**. All cell lines were successfully infected with SARS-CoV-2 (**Figure 5B, Table S2D**), but *ACE2* expression was not affected by infection in any cell line tested (**Figure 5A, Table S2D**). Induction of *dACE2* expression tracked with previously reported SARS-CoV-2 infectivity rates in these cells^23^. Specifically, *dACE2* was most strongly induced in Caco-2 cells, in which over 80% of cells were infected by 24 hrs, moderately induced in T84 cells (20% of infected cells), and not induced in Calu3 cells (10% of infected cells). A similar expression pattern was observed for an ISG *IFIT1* except for Calu3 cells, in which *IFIT1* was significantly induced (**Figure 5A, Table S2D**). We performed a similar analysis in human colon and ileum organoid cultures derived from two donors. There was more variability in expression of the three genes tested but, overall, the expression of *dACE2* and *IFIT1* but not *ACE2* was significantly induced by SARS-CoV-2 infection (**Figure 5C, Table S2E**), despite comparable SARS-CoV-2 infection load (**Figure 5D, Table S2E**), and a similar infection rate (10% of the cells)^23^. These results confirm that *dACE2* is inducible by SARS-CoV-2 infection.

**Figure 5.**
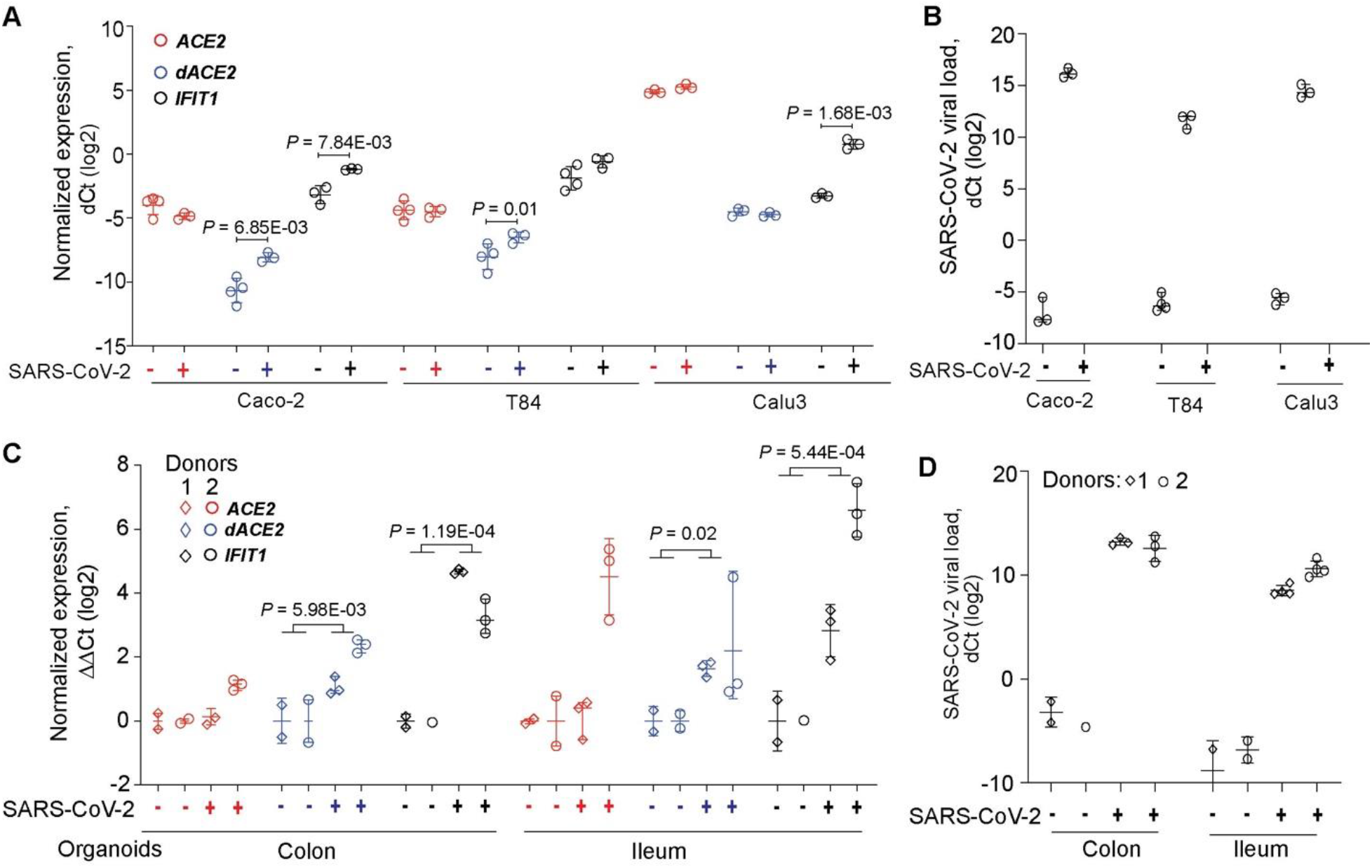
*dACE2* is induced by SARS-CoV-2 in human cell lines and organoid cultures. Expression of *ACE2*, *dACE2* and a control ISG *IFIT1* in **A)** colon cancer cell lines Caco2 and T84 and a lung cancer cell line Calu3 and **C)** colon and ileum organoid cultures derived from two donors; **B** and **D)** SARS-CoV-2 viral loads in corresponding cells. P-values are for Student’s T-test.

### dACE2 is non-functional as SARS-CoV-2 receptor and carboxypeptidase

Despite the strong induction of *dACE2* mRNA expression, we were unable to detect endogenous dACE2 in SeV-infected cell lines by Western blotting with commercial ACE2 antibodies (data not shown). However, in the proteome database of mass spectrometry data available for breast, colon, and ovarian TCGA tumors^25^, we detected human-specific peptides matching the 10 aa encoded by the *dACE2*-Ex1c (**Figure S10**), suggesting that dACE2 protein could be expressed in some conditions.

The substantial N-terminal truncation by 356 aa in the peptidase domain of the putative dACE2 protein is expected to have significant functional consequences compared to the activity of the full-length ACE2 protein of 805 aa. For example, decreased or no binding by the SARS-CoV-2 spike receptor-binding domain (RBD) would be expected for dACE2. Indeed, when we treated cells overexpressing GFP or Myc-DDK-tagged dACE2 or ACE2 with biotinylated spike protein RBD (**Figure 6A**), dACE2 failed to bind and internalize the RBD (**Figure 6B, C**) or affect the RBD binding by ACE2 (**Figure 6D-F**). The N-terminal truncation is also predicted to affect carboxypeptidase activity of dACE2, which is important for its ability to cleave angiotensin II, des-Arg9-bradykinin, and other substrates of ACE2. Indeed, we observed carboxypeptidase activity in lysates of cells transfected with ACE2-GFP but not dACE2-GFP (**Figure 6G, H**). Thus, the main activities that involve the peptidase domain of ACE2 appear to be abrogated in dACE2.

**Figure 6.**
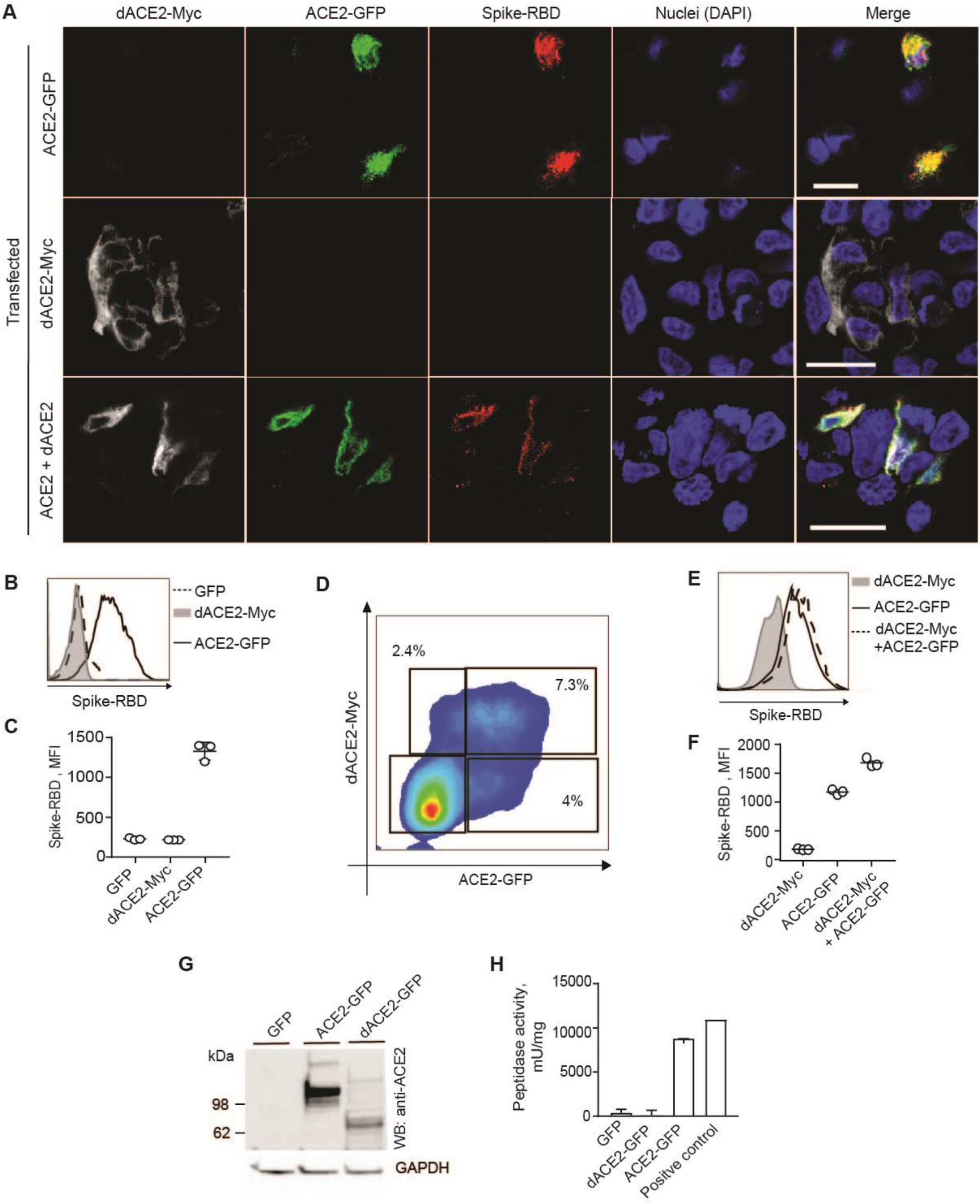
dACE2 is non-functional for binding SARS-CoV-2 spike protein RBD and as a carboxypeptidase. **A)** Representative confocal images of T24 cells transiently overexpressing dACE2-Myc (white), ACE2-GFP (green) and treated with receptor-binding domain (RBD) of SARS-CoV-2 spike protein (red), nuclei (DAPI)-blue; bars=20μM. **B-D)** Representative flow cytometry histogram **B)** and mean fluorescence intensity (MFI) values from 3 biological replicates **C)** of spike-RBD binding to the surface of ACE2-GFP but not dACE2-Myc expressing T24 cells. **D)** Representative flow cytometry plot showing gates for T24 cells co-transfected with dACE2-Myc and ACE2-GFP. Gates were drawn to identify cells expressing dACE2-Myc, ACE2-GFP, or both proteins. **E)** Histogram and **F)** plot depicting MFI of spike protein-RBD binding in gated cells from plot **E**, based on 3 biological replicates. Data represent one of two independent experiments. **G)** A representative Western blot showing detection of ACE2-GFP and dACE-GFP in protein lysates of T24 cells transfected with equal amounts of plasmids. Based on Western blot densitometry, equal estimated amounts of ACE2 and dACE2 proteins were used in peptidase activity assays in **H**. Results are based on 3 biological replicates.

## DISCUSSION

*ACE2* was recently proposed to be an ISG due to its induction in IFN-high conditions, raising concerns about its potential role in increasing SARS-CoV-2 infection^4,5^ and the safety of IFN-based treatments proposed for COVID-19. Our discovery of *dACE2*, a primate-specific version of *ACE2,* demonstrates that it is *dACE2* and not *ACE2* that is induced by IFNs and viruses, including SARS-CoV-2. dACE2, however, did not appear to bind SARS-CoV-2 spike protein or affect the binding of ACE2 in our *in vitro* experiments, thus assuring that ISG-type induction of *dACE2* would not increase viral entry.

Our findings highlight that in the presence of *dACE2*, *ACE2* expression is likely to be misinterpreted when detected by methods such as 3’-scRNA-seq and gene-based assays that target shared *dACE2* and *ACE2* sequences. However, in non-primate species or in human cells that do not express *dACE2*, the same detection methods would correctly quantify *Ace2 (ACE2),* but not finding it is an ISG. These considerations are also important when using non-primate species to study SARS-CoV-2 infection and COVID-19 because any possible effects contributed by *dACE2* will not be captured in these animal models.

By analyses in multiple human cell types and tissues, we showed that expression of *dACE2,* but not *ACE2,* is inducible by IFNs (type I, II and III) and viruses that induce IFN responses. Suppression of IFN signaling by SARS-CoV-2 has been reported by several studies^26,27^, possibly explaining only a moderate effect of SARS-CoV-2 infection on *dACE2* induction in our experiments. While type I and type III IFNs are barely detectable in COVID-19 patients^28,29^, high levels of IFN-γ have been reported^29,30^. Specifically, a 3’-scRNA-seq analysis showed *ACE2* induction by SARS-CoV-2 infection in ciliated epithelia, where high levels of IFN-γ producing immune cells were also detected^5^. Our results strongly suggest that the induction of *dACE2* and not *ACE2* was detected in these COVID-19 patients.

IFN-γ-driven ISG signatures are contributed by immune cells that infiltrate tumors and inflamed virally infected tissues. We could not study the involvement of *dACE2* in these signatures in cell culture models because they lack the immune component; in tissues by 3’-scRNA-seq because this method does not differentiate *ACE2* and *dACE2*; and in mice – because they lack *dAce2*. Instead, we explored the extensive TCGA dataset of more than 10,000 tumors in which we *de-novo* quantified *dACE2* expression based on RNA-seq data. We believe the conclusions of these analyses can be extended to inflamed virally infected tissues, for which comparable RNA-seq data is limited by small sample sets; low percentage of mappable reads due to substantially degraded input RNA, or raw data is not provided due to patient privacy restrictions.

Although all IFNs (type I, II and III) appeared to be potent inducers of *dACE2* expression *in vitro*, IFN-γ might be an important driver of *dACE2* expression in tissues with significant infiltration by immune cells, such as in tumors or virally infected tissues. Furthermore, the expression patterns observed in TCGA indicated that *dACE2* expression might be intrinsically enriched in squamous epithelial cells, which give rise to corresponding tumors of the respiratory, gastrointestinal, and urogenital tracts.

While our experiments did not yet identify a clear functional role for *dACE2*, the absence in non-primate species and the high degree of *dACE2*-Ex1c sequence conservation over 43 million years of primate evolution^31^ argues for a potentially important role of *dACE2.* Endogenous dACE2 was not detected in cell lines by Western blotting with anti-ACE2 antibodies and may require the development of dACE2-specific antibodies. However, the detection of dACE2-specific peptides in some TCGA tumors suggests that the protein production occurs, but might require some specific conditions or better detection tools. However, based on our *in vitro* data, it is unlikely that dACE2 protein would affect the binding and cellular access of SARS-CoV-2.

Although possible ISG-type ACE2 induction was considered as a risk for increasing SARS-CoV-2 infection, ACE2 deficiency rather than overexpression is discussed as a major factor contributing to COVID-19 morbidity^12–14,32^. Functional ACE2 deficiency occurs due to internalization of the SARS-CoV-2 -ACE2 complex^2,33^, which restricts ACE2 from performing its physiological functions, including its role as carboxypeptidase for angiotensin II and des-Arg9-bradykinin and other peptide hormones. ACE2 deficiency might also be created due to its regulation at the mRNA level, which needs to be explored.

Cancer patients are considered to be at a higher risk of more severe COVID-19 outcomes compared to the general population^34,35^ due to older age, comorbidities and effects of cancer and cancer treatments. Lung cancer patients are at a specifically increased risk of severe COVID-19 outcomes^34^. In our analysis, *dACE2* expression was common in tumors and particularly enriched in lung tumors of bronchial origin (LUSC), where the proper function of ACE2 is essential for protection from virus-induced tissue damage. The possible role of *dACE2* expression and COVID-19 outcomes, specifically in cancer patients, should be further explored. ACE inhibitors (ACEIs) and angiotensin-receptor blockers (ARBs) are widely used to control hypertension and treat heart disease and chronic kidney disease^10^. Significant concerns were raised that ACEIs and ARBs could induce ACE2 expression, leading to increased SARS-CoV-2 infection and possibly accounting for COVID-19 severity and high mortality in those who are likely to use these medications - the elderly and patients with cardiovascular disease. We demonstrated that *ACE2* expression is not inducible by IFNs, but it would be important to explore the effects of ACEIs and ARBs on *dACE2* expression to properly assess this risk. The effects of other factors, such as smoking on *dACE2* expression, should also be considered.

In conclusion, we present the first report of the discovery and functional annotation of *dACE2*, an IFN-inducible isoform of *ACE2*. The existence of two functionally distinct *ACE2* isoforms reconciles several biological properties previously attributed to ACE2, with *dACE2* being an ISG, and ACE2 acting as the SARS-CoV-2 entry receptor and carboxypeptidase, without being regulated by IFNs. While our understanding of the functional role of *dACE2* is still unfolding, we believe these insights will clarify our knowledge on ACE2 and provide new research leads in understanding COVID-19 susceptibility, mechanisms, and outcomes.

## MATERIALS AND METHODS

### Cells

All cell lines and primary cells used are listed in **Table S3**. Cell lines were either used within 6 months after purchase or were periodically authenticated by microsatellite fingerprinting (AmpFLSTR Identifiler, Thermo Fisher) by the Cancer Genomics Research Laboratory/NCI). All cell lines were regularly tested for mycoplasma contamination using the MycoAlert Mycoplasma Detection kit (Lonza). The previously described^22^ normal human bronchial epithelial cells (NHBE) were isolated from normal lungs that were not used for transplantation. The lungs were obtained from 5 de-identified donors via a tissue retrieval service (International Institute for the Advancement of Medicine, Edison, NJ) with ethical approval from the Conjoint Health Research Ethics Board of the University of Calgary and the Internal Ethics Board of the International Institute for the Advancement of Medicine. Anonymized human tissue from colon resections was obtained from the University Hospital Heidelberg, in accordance with the recommendations of the University Hospital Heidelberg and written informed consent obtained from all subjects in accordance with the Declaration of Helsinki. The protocol (S-443/2017) was approved by the Ethics Commission of the University Hospital Heidelberg. Organoids were generated from these tissues as previously described^23^.

### Viral infections

Stocks of Sendai virus (SeV) Cantell strain were purchased from Charles River Laboratories. Cells listed in **Table S3** were infected in duplicates or triplicates with SeV (7.5×10^5^ CEID50/ml) for 12 or 24 hrs as previously described^19–21^.

Infections with SARS-CoV-2 in colon cancer cell lines (Caco2 and T84) and lung cancer cell line (Calu3) and colon and ileum organoid cultures were previously described^23^. Briefly, the infections were performed using the MOI=1 for cell lines. Organoids were infected with 3×10^5^ FFU of the virus, based on titers in Vero E6 cells. Culture media was removed from cells/organoids and the virus was added for 1 hr at 37°C. After virus removal, cells were washed 1x with PBS, and media was added back to the cells. Viral replication and ISG induction were monitored for 24 hrs post-infection. The percentage of infected cells was determined by immunofluorescence staining and quantification of the number of nuclei vs. the number of infected cells. It was determined that infection rates were 80% in Caco2 cells, ~20% in T84 cells, ~10% in Calu3 cells, and 10% in organoids.

### PCR, cloning and Sanger sequencing

cDNA was synthesized from 250 ng of total RNA per 20 μl reactions using the RT2 First-Strand cDNA kit and random hexamers (Qiagen). PCR for the full-length *dACE2* was performed using primers and conditions listed in **Table S4**. PCR-amplified products were resolved on 1% agarose gel, cut, purified, and Sanger-sequenced. After cDNA sequences were validated, constructs for dACE2 with C-terminal Myc-DDK and GFP tags cloned in pcDNA3.4 vector were custom-synthesized by Thermo Fisher. ACE2 with C-terminal GFP-tag (RG208442) and Myc-DDK tag (RC208442) were purchased from Origene. Empty vectors pMax-GFP (Lonza) and pCMV6-AC-Myc-DDK (Origene) were used as controls.

### Treatments with IFNs

All IFNs used are listed in **Table S5**. The IFN-treated NHBE cells were previously described^22^. Briefly, the cells were cultured in BEGM with supplements (Lonza), plated in 6-well plates and utilized at ~70% confluence (typically after 10–11 days with media change every 2 days). Cells were left untreated or treated with IFN-α2b (INTRON A, Merck, 100 IU/ml) or IFN-λ3 (R&D Systems, 100ng/ml) for 24 hrs. Cells were washed with PBS, resuspended in TRIzol (Thermo Fisher) and stored at −80°C for future RNA isolation. Total RNA was extracted with Direct-zol mini RNA isolation kits (Zymo Research). The IFN treatment of human colon and ileum organoids was previously described^23^. Briefly, cells were seeded 24 hrs prior to IFN treatment to reach a confluency of 70% at the time of treatment. Media was removed from cells and replaced with a cocktail of IFN-λ1-3 (100ng/mL of each for a total of 300ng/mL). Media was removed 24 hrs post-treatment, and RNA was extracted using the RNAeasy kit (Qiagen). 250ng of RNA was used to prepare cDNA using iSCRIPT (BioRad) and qRT-PCR was performed using iTaq SYBR Green (BioRad). T84 and Caco-2 cells were treated with IFN-γ (2 ng/ml) for 48 hrs, followed by cell harvesting, RNA extraction and expression analysis.

### qRT-PCR

cDNA was synthesized from the total RNA with the RT^2^ First Strand Kit (Qiagen), Superscript VILO IV (Thermo Fisher), or iSCRIPT cDNA kit (BioRad), always with an additional DNase I treatment step. qRT-PCR assays were performed in technical duplicates in 96- or 384-well plates on QuantStudio 7 (Life Technologies) or Bio-Rad CFX 96 instrument, with RT² SYBR Green (Qiagen), POWER SYBR (Thermo Fisher), iTaq SYBR (BioRad) or TaqMan (Thermo Fisher) expression assays (**Table S4**). The expression of target genes was normalized by geometric means of endogenous controls (*GAPDH, HPRT1, TBP* or *ACTB,* as indicated in **Table S2**), and presented as dCt values relative to endogenous controls (log2 scale).

### RNA sequencing (RNA-seq) of T47D and RT4 cells

Total RNA was extracted from T47D and RT4 cells using RNeasy Mini Kit with an on-column DNase digestion (Qiagen) and treated with Ribo-Zero (Illumina). RNA-seq libraries were prepared from high-quality RNA samples (RIN scores >9.0) with KAPA Stranded RNA-seq Kit with RiboErase (Roche). Paired 150-bp reads (21.2 - 118.8 million reads per sample) were generated with HiSeq 2500 (Illumina) by the Cancer Genomics Laboratory (DCEG/NCI). The reads were aligned with STAR alignment tool version 7.1.2a (21) using the GRChg37/hg19 genome assembly and visualized using the Integrative Genomics Viewer (IGV).

### RNA-seq analysis of data from NCBI Sequence Read Archive (SRA) and TCGA

RNA-seq datasets (**Table S6**) were downloaded from NCBI SRA using SRA tools. The FASTQ files were compressed using GZIP and aligned with STAR version 7.1.3a to the GRChg38/hg38 genome assembly. BAM files with less than 80% of mappable reads were excluded. BAM files were indexed and sliced using SAM tools to include 51.6 Kb of the *ACE2* genomic region (chrX:15,556,393-15,608,016, hg38). For mouse RNA-seq data sets, the alignment was done with reference genome mm10. For TCGA STAR-aligned RNA-seq data, BAM slices for the *ACE2* region were acquired for 11, 041 TCGA samples (10,328 tumors and 713 tumor-adjacent normal tissues) through the NCI Genomics Data Commons (GDC) portal accessed on May 12, 2020, using workflow https://docs.gdc.cancer.gov/API/Users_Guide/BAM_Slicing/.

### Estimation of RNA-seq read counts for *ACE2* exons

RNA-seq reads corresponding to the three first exons, *ACE2*-Ex1a and Ex1b and *dACE2-*Ex1c, were counted by processing RNA-seq BAM slices using R package ASpli version 1.5.1 with default settings. Genomic coordinates were manually curated in the GTF file and the ‘counts’ function was used to generate and export RNA-seq reads for the selected exons in a tab file format. The reads were normalized by dividing by the exon length (**Table S7**) and multiplying by the geometric mean of the total reads of the three exons (Ex1a, Ex1b and Ex1c) across all samples as a scaling factor to adjust for variability in sequencing coverage between samples. The analysis of exon expression patterns within tissue subtypes was based on log 2-transformed raw reads for each exon. Correlation analysis of normalized expression of *ACE2-*Ex1a and Ex1b and *dACE2-*Ex1c with TPMs of IFNs, STATs, IRFs and select ISG controls (*ISG15* and *ISG20*) was performed in R using package dplyr.

Expression values for all *IFN* genes for all tumors in TCGA PanCancer Atlas were downloaded as RSEM values from cBioPortal for Cancer Genomics (https://www.cbioportal.org/). Expression of *IFNL4*, a most recently discovered IFN^36^, was not available in TCGA dataset based on hg19/GRCh37 reference. Because *IFNG* was expressed in most samples compared to other IFN genes (**Figure S8A**), it was used for further analysis. Correlation analyses of the normalized expression of *ACE2-*Ex1b and *dACE2-*Ex1c were performed with log2(RSEM+1) values for *IFNG*. Correlation patterns between *IFNG* and *ACE2*-Ex1a were similar to those between *IFNG* and *ACE2*-Ex1b (data not shown).

### Unsupervised clustering and correlation analyses in TCGA

Gene expression Z-scores in the lung squamous cell carcinoma (TCGA-LUSC, n=501) dataset were calculated for 270 ISGs from a previously curated list^24^. ISGs with low expression values, (below 10 reads), or expressed in less than 5% of tumors were excluded. The data was used for self-organizing maps (SOM) clustering, which is an unsupervised machine learning approach enabling data dimensionality reduction without relying on any assumption about the data structure^37,38^. The SOM algorithm was iterated 100,000 times with Euclidean distance, linearly decreasing the learning rate from 0.05 to 0.01 using the “Kohonen” R package. The ISG expression patterns were projected onto a two-dimensional 10×10 hexagonal map. Thus, each node in this map is an expression profile representing a subset of the samples. SOM output, trained based on 100,000 iterations, was used to estimate the contribution of each ISG to defining the clusters as a variance weighted according to the size of each node. A total of 6 clusters were estimated by kmeans algorithm and used to generate an expression heatmap. Expression Z-scores of the top 100 ISGs ranked based on their contribution to defining the clusters were plotted using the “pheatmap” R package. Pearson correlation coefficients and corresponding FDR-adjusted p-values were calculated between Z-scores for the top 100 ISGs and both *ACE2* and *dACE2* in cluster 5, which included 114 tumors with the highest ISG expression. The analysis was performed using the “Hmisc” R package.

### In *silico* analysis of promoter regulatory elements relevant for IFN signaling

Promoters were defined within the −800 bp/+100 bp window from the corresponding transcription start sites (TSS). The window was limited by 800 bp, based on the intronic distance between TSS of Ex1c and its upstream exon. Promoters of *ACE2*-Ex1a (P1) and Ex1b (P2) and *dACE2-*Ex1c (P3) were analyzed using Nsite tool^39^ from online bioinformatics gateway Softberry (www. softberry.com) to predict transcription factor binding sites. The search was set against the ooTFD (Object-oriented Transcription Factors Database) of human and animal transcription factor binding sites largely curated according to the functional data from the literature. The parameters were set to allow a maximum of 1 or 2 mismatches with the known motifs. ISG-type motifs were manually curated from ~300 predicted and annotated motifs per promoter.

### Mining of proteomics datasets

Mass-spectrometry datasets generated for TCGA colon, breast and ovarian tumors (http://www.pepquery.org/) were mined for matches to the 36-aa fragment of dACE2, including the unique 10 aa encoded by *dACE2*-Ex1c. The analysis was done with PepQuery peptide-centric search engine^25^, using the following parameters: MS dataset of a specific cancer type; target event as protein; scoring algorithm as Hyperscore and not selecting for Unrestricted modification filtering. The identified peptides for each cancer type were exported as CSV files and manually analyzed for further assessment of peptide quality.

### Transient transfections

Transient transfections were performed with Lipofectamine 3000 (Thermo Fisher). Unless specified, bladder cancer cells (T24), in which no baseline expression of *ACE2* or *dACE2* was detected (**Table S2**), were used for transfections at 70–90% confluency in 12- or 6-well plates for 24 hrs.

### Western blot

Cells were lysed with RIPA buffer (Sigma) supplemented with protease inhibitor cocktail (Promega) and PhosSTOP (Roche) and placed on ice for 30 min, with vortexing every 10 min. Lysates were pulse-sonicated for 30 sec, with 10-s burst-cooling cycles, at 4°C, boiled in reducing sample buffer for 5 min and resolved on 4–12% Bis-Tris Bolt gels and transferred using an iBlot 2 (Thermo Fisher). Blots were blocked in 2.5% milk in 1% TBS-Tween before staining with antibodies (**Table S5**). The signals were detected with HyGLO Quick Spray (Denville Scientific) or SuperSignal West Femto Maximum Sensitivity Substrate (Thermo Fisher) and viewed on a ChemiDoc Touch Imager with Image Lab 5.2 software (BioRad).

### Confocal microscopy

T24 cells were transiently transfected with ACE2-GFP, dACE2-Myc-DDK, or co-transfected with both constructs in 4-well chambered slides (2×10^4^ cells/well, LabTek). After 24 hrs, cells were treated with 2 ng/ml of recombinant biotinylated SARS-CoV-2 spike protein RBD (spike protein-RBD, Sino Biological) for 1 hour at 37°C. Cells were washed twice with media and then stained with 5μg/ml streptavidin PE (Thermo Fisher) for 30 min at 37°C. Cells were then washed twice with PBS and fixed with 4% paraformaldehyde (BD Biosciences) for 30 min. After rinsing twice in PBS and permeabilization buffer (BD Biosciences), cells were incubated with permeabilization buffer for 1hr. Fixed cells were incubated with rabbit anti-FLAG antibody (1:250 dilution, Thermo Fisher) overnight, washed and then stained with anti-rabbit Alexa Fluor 680 (1:500 dilution, Thermo Fisher). Slides were mounted with antifade mounting media with DAPI (Thermo Fisher) and imaged at 40X magnification on an LSM700 confocal laser scanning microscope (Carl Zeiss) using an inverted oil lens.

### Flow cytometry analysis of SARS-CoV-2 spike protein-RBD binding

T24 cells were transiently transfected with ACE2-GFP, dACE2-Myc-DDK, or co-transfected with both constructs in 12-well plates (1×10^5^ cells/well). After 24 hrs, cells were stained with recombinant biotinylated spike protein RBD as described above and analyzed with multiparametric flow cytometry on a FACS Aria III (BD Biosciences) and FlowJo.v10 software (BD Biosciences).

### Peptidase activity assays

T24 cells were transiently transfected with ACE2-GFP, dACE2-GFP, or empty GFP-vector in triplicates. After 24 hrs, cells were pelleted and lysed with 400 μl of Lysis Buffer provided with the ACE2 activity kit (#K897, BioVision). Cell lysates were analyzed by Western blots with C-terminal anti-ACE2 antibody (Abcam), and the amounts of recombinant ACE2-GFP and dACE2-GFP proteins in lysates were quantified by densitometry analysis (Imagelab, BD Biosciences) of Western blots. The amounts of ACE2 and dACE2 lysates to be used in protease assays were determined based on protein quantification. Protein lysates were processed in duplicates using the kit reagents and according to protocol. The activity was measured as fluorescence (Ex/Em = 365/410-460 nm) using Promega GlowMax plate reader for two time points between 30 mins and 2 hrs after adding the corresponding substrate mix and normalizing for baseline level. Positive control was provided by the kit.

### Statistical analysis

Expression of *ACE2*-Ex1a and Ex1b and *dACE2*-Ex1c between groups of samples was evaluated by the non-parametric Mann–Whitney U test. Student’s T-test was used for small groups of samples. Statistical tests for other analyses are indicated in corresponding sections. FDR adjustment was applied when indicated. P-values <0.05 were considered significant.

### Computational resources

We used the NIH Biowulf supercomputing cluster and specific packages for R version 3.6.2.

## Acknowledgments

We thank Nathan Cole (DCEG/NCI) for help with the acquisition of TCGA sliced BAM files and Dr. David Proud (University of Calgary) for providing NHBE cells. We thank the CGR/DCEG/NCI for help with RNA-sequencing and authentication of cell lines by Identifiler profiling. The results are partially based on data generated by TCGA Research Network. The project was supported by the Intramural Research Program of the Division of Cancer Epidemiology and Genetics, National Cancer Institute; HP was supported by the NIH grant R21AG064479-01 and a Brain Health Research Institute Pilot Award from Kent State University; SB and MS received financial support from the Deutsche Forschungsgemeinschaft (DFG) project number 240245660 (SFB1129) 415089553 (Heisenberg) and 272983813 (TRR179) to SB and project number 416072091 to MS. DMS and DLJT received financial support from the Li Ka Shing Institute of Virology.

NCBI GenBank accession number for dACE2: MT505392

URLs:

UCSC Genome Browser: https://genome.ucsc.edu/

TCGA Research Network: https://www.cancer.gov/tcga

## Conflict of interest statement

Authors declare no competing interests

## Notes

### Competing Interest Statement

The authors have declared no competing interest.

