## Supplementary Materials for "Interferons and viruses induce a novel primate-specific isoform dACE2 and not the SARS-CoV-2 receptor ACE2"

Onabajo, Banday et al,

### Supplementary Tables

**Table S1.** Expression of IFNs and select ISGs in T47D cells at baseline and after SeV infection, RNA-seq FPKM, hg38

| Gene | IFN type | Untreated 1 | Untreated 2 | SeV 12hrs.1 | SeV 12hrs.2 |
|---|---|---|---|---|---|
| IFNA1 | Type I | 0.00 | 0.00 | 0.52 | 0.27 |
| IFNA2 | Type I | 0.00 | 0.00 | 0.13 | 0.04 |
| IFNA4 | Type I | 0.00 | 0.00 | 0.00 | 0.00 |
| IFNA5 | Type I | 0.00 | 0.00 | 0.00 | 0.00 |
| IFNA6 | Type I | 0.00 | 0.00 | 0.00 | 0.00 |
| IFNA7 | Type I | 0.00 | 0.00 | 0.30 | 0.25 |
| IFNA8 | Type I | 0.00 | 0.00 | 0.00 | 0.05 |
| IFNA10 | Type I | 0.00 | 0.00 | 0.24 | 0.39 |
| IFNA13 | Type I | 0.00 | 0.00 | 0.33 | 0.09 |
| IFNA14 | Type I | 0.00 | 0.00 | 0.00 | 0.08 |
| IFNA16 | Type I | 0.00 | 0.00 | 0.00 | 0.11 |
| IFNA17 | Type I | 0.00 | 0.00 | 0.00 | 0.00 |
| IFNA21 | Type I | 0.00 | 0.00 | 0.00 | 0.05 |
| **IFNB1** | **Type I** | **0.00** | **0.00** | **337.30** | **381.62** |
| IFNE | Type I | 0.00 | 0.00 | 0.04 | 0.03 |
| IFNK | Type I | 0.00 | 0.00 | 0.00 | 0.00 |
| IFNG | Type II | 0.00 | 0.00 | 0.00 | 0.00 |
| **IFNL1** | **Type III** | **0.00** | **0.00** | **252.53** | **292.93** |
| **IFNL2** | **Type III** | **0.00** | **0.00** | **110.41** | **123.48** |
| **IFNL3** | **Type III** | **0.00** | **0.00** | **122.21** | **138.66** |
| **IFNL4** | **Type III** | **0.00** | **0.00** | **122.77** | **132.11** |
| ISG15 | ISG | 4.29 | 3.25 | 2698.66 | 3206.30 |
| IFIT1 | ISG | 0.94 | 0.81 | 879.53 | 1033.89 |
| MX1 | ISG | 5.20 | 4.66 | 849.95 | 881.69 |

FPKM - fragments per kilobase of exon per million reads mapped

**Table S2. Expression of *ACE2* and *dACE2* in various cell lines and conditions** (separate Excel file)

**Table S3. Cell lines used**

| **Cells** | **Cell type** | **Source** | **Media** |
|---|---|---|---|
| Primary tonsil epithelial cells | Normal tissue from donors | ScienCell | Tonsil Epithelial Cell Medium |
| T47D (MDA-MB-23) | Breast cancer | ATCC | DMEM |
| T24 | Bladder cancer | ATCC | McCoy's 5A |
| HT-1376 | Bladder cancer | ATCC | DMEM |
| HTB-9 | Bladder cancer | ATCC | RPMI-1640 |
| RT-4 | Bladder cancer | ATCC | McCoy's 5A |
| HBLAK | Immortalized uroepithelial | CELLnTEC | CnT-Prime |
| PC3 | Prostate cancer | ATCC | F-12 |
| 22RV1 | Prostate cancer | ATCC | RPMI |
| DU145 | Prostate cancer | ATCC | EMEM |
| HepG2 | Liver cancer | ATCC | DMEM |
| Caco-2 | Colon cancer | ATCC | EMEM |
| T84 | Colon cancer | ATCC | DMEM: F-12 |
| A549 | Lung cancer | ATCC | F-12 |
| Calu3 | Lung cancer | ATCC | DMEM |
| Capan-1 | Pancreatic cancer | ATCC | IMDM |
| HeLa | Cervical cancer | ATCC | EMEM |
| TCCSUP/HTB5 | Bladder Cancer | ATCC | EMEM |
| 5637/HTB9 | Bladder Cancer | ATCC | RPMI |
| J82 | Bladder Cancer | ATCC | EMEM |
| SW780 | Bladder Cancer | ATCC | Leibovitz's L-15 Medium |
| UMUC3 | Bladder Cancer | ATCC | EMEM |
| 293T | Kidney | ATCC | DMEM |
| NHBE | Primary normal human bronchial epithelial cells from 5 donors Described in (Santer et al., 2020) | International Institute for the Advancement of Medicine | BEGM (bronchial epithelial growth medium) + bulletkit |
| Organoid cultures of colon and ileum | Described in (Stanifer et al., 2020) | University Hospital Heidelberg | Human organoid media |

ATCC - American Type Culture Collection

**Table S4. Primers and expression assays used**

| Primers | Sequence | Assay type, amplicon size |
|---|---|---|
| ACE2_F | GGGCGACTTCAGGATCCTTAT | ACE2 SYBR Green assay, 80 bp |
| ACE2_R | GGATATGCCCCATCTCATGATG | |
| dACE2_F | GGAAGCAGGCTGGGACAAA | dACE2 SYBR Green assay, 73 bp |
| dACE2_R | AGCTGTCAGGAAGTCGTCCATT | |
| ACE2_F | GGGCGACTTCAGGATCCTTAT | ACE2 TaqMan assay, 80 bp |
| ACE2_R | GGATATGCCCCATCTCATGATG | |
| ACE2_probe | ATGGACGACTTCCTGACAG | |
| dACE2_F | GGAAGCAGGCTGGGACAAA | dACE2 TaqMan assay, 73 bp |
| dACE2_R | AGCTGTCAGGAAGTCGTCCATT | |
| dACE_probe | AGGGAGGATCCTTATGTG | |
| dACE2_F | AGTGCTTCATTGAGGAGAGCTCT | dACE2, 5'-3'UTR, 1535 bp |
| dACE2_R | TCTATACCATGAAATTAACATTTACATACAAC | 98°C-30s, 98°C-10s, 60°C-30s, 72°C-40s, 35 cycles, 72°C-2 min; Q5 High-Fidelity 2X PCR Master Mix (NEB) |
| HPRT1_F | TGACACTGGCAAAACAATGCA | SYBR Green assay, 94 bp |
| HPRT1_R | GGTCCTTTTCACCAGCAAGCT | |
| MX1_F | ACCTGATGGCCTATCACCAG | SYBR Green assay, 154 bp |
| MX1_R | TTCAGGAGCCAGCTGTAGGT | |
| IFIT1_F | AAAAGCCCACATTTGAGGTG | SYBR Green assay |
| IFIT1_R | GAAATTCCTGAAACCGACCA | SYBR Green assay |
| GAPDH | Hs04420632_g1 (Thermo Fisher) | TaqMan assay |
| ACTB | 4352667 (Thermo Fisher) | |
| ISG15 | Hs01921425_s1 (Thermo Fisher) | TaqMan assay |

**Table S5. Reagents used**

| Antibodies | | | | | | |
|---|---|---|---|---|---|---|
| **Target gene** | **Cat. No.** | **Source** | **Target species** | **Host** | **Tag** | **Dilution** |
| ACE2 | ab15348 | Abcam | Human | Rabbit | | 1:250 |
| Myc-DDK | | Thermo Fisher | Tag | Rabbit | | 1:1000 |
| GAPDH | Ab9485 | Abcam | Human | Rabbit | | 1:1000 |
| GFP | MA515256 | Thermo Fisher | Tag | Mouse | | 1:1000 |
| DYKDDDDK Epitope Tag | NB600-347 | Novus Biologicals | Tag | Goat | | 1:1000 |
| IgG | #7074 | Cell Signaling | Rabbit | Goat | HRP | 1:5000 |
| IgG | sc2314 | Santa Cruz | Mouse | Donkey | HRP | 1:5000 |
| IgG | sc2304 | Santa Cruz | Goat | Donkey | HRP | 1:5000 |
| Streptavidin | SA10044 | Thermo Fisher | Tag | | PE | 1:200 |
| IgG | A32734 | Thermo Fisher | Rabbit | Goat | AF680 | 1:200 |

| Interferons | | | | |
|---|---|---|---|---|
| **IFN** | **Source** | **Concentration** | **Time** | **Experiment** |
| IFNa2b | Merck, Intron A | 100 IU/ml | 24 hrs | NHBE |
| IFN-λ3 | R&D Systems, Cat# 5259-IL/CF | 100 ng/ml | 24 hrs | NHBE |
| IFN-β1 | Biomol, Cat#86421 | 2000IU/mL | 24 hrs | Organoids Cell lines |
| IFN-λ1 | Peprotech, Cat#300-02L | 100 ng/ml | 24 hrs | Organoids Cell lines |
| IFN-λ2 | Peprotech, Cat#300-02K | 100 ng/ml | 24 hrs | Organoids Cell lines |
| IFN-λ3 | Biomol, Cat#179-ML-025 | 100 ng/ml | 24 hrs | Organoids Cell lines |
| IFN-β | R&D Systems, Cat# 8499-IF | 0.5 ng/ml | 48 hrs | Cell lines |
| IFN-γ | R&D Systems Cat# 285-IF | 2 ng/ml | 48 hrs | Cell lines |

**Table S6. RNA-seq datasets analyzed**

| Datasets | NCBI SRA | Alignment reference genome | Reference |
|---|---|---|---|
| Breast cancer cell line T47D, SeV-infected for 12 hours (n=2), not infected, n=2 | PRJNA512015 | hg19 | Current work |
| Nasal epithelial cells from 30 asthmatic patients were infected with rhinovirus strains - RV-A16 (n=30), RV-C15 (n=30) or not infected (n=30) | PRJNA627860 | hg38 | NA |
| Human lung explants infected with influenza A/H3N2 virus from 5 donors, n=20 | PRJNA557257 | hg38 | NA |
| Lung cells infected with the respiratory syncytial virus (RSV): human lung mucoepidermoid pulmonary carcinoma cell line H292, RSV-infected (n=1) and mock (n=1); lung cells from mice infected with RSV, n=3 and mock, n=3 | PRJNA588982 | hg38 and mm10 | (McAllister et al., 2020) |
| Normal human tissues, n = 95 | PRJEB4337 | hg38 | (Fagerberg et al., 2014) |

**Table S7. Nucleotide sequences and genome coordinates of three alternative first exons of *ACE2* and *dACE2* used for quantification of RNA-seq reads**

| Exon | Sequence | Coordinates, hg38 | Length, bp | RefSeq ID |
|---|---|---|---|---|
| ACE2, Ex1a | GGCACTCATACATACACTCTGGCA<br>ATGAGGACACTGAGCTCGCTTCTG<br>AAATTTGACAAGATAACCACTAAA<br>ATCTCTTTGAATTCTATGTTGTTGT<br>GATCCCATGGCTACAGAGGATCAG<br>GAGTTGACATAGATACTCTTTGGAT<br>TTCATACCATGTGGAGGCTTTCTTA<br>CTTCCACGTGACCTTGACTGAGTTT<br>TGAATAG | chrX:15,601,956-15,602,158 | 203 bp | NM_021804.3 |
| ACE2, Ex1b | CGCCCAACCCAAGTTCAAAGGCTG<br>ATAAGAGAGAAAATCTCATGAGGA<br>GGTTTTAGTCTAGGGAAAGTCATTC<br>AGTGGATGTGATCTTGGCTCACAG<br>GGGACGATGTCAAGCTCTTCCTGG<br>CTCCTTCTCAGCCTTGTTGCTGTAA<br>CTGCTGCTCAGTCCACCATTGAGG<br>AACAGGCCAAGACATTTTTGGACA<br>AGTTTAACCACGAAGCCGAAGACC<br>TGTTCTATCAAAGTTCACTTGCTTC<br>TTGGAATTATAACACCAATATTACT<br>GAAGAGAATGTCCAAAACATG | chrX:15,600,726-15,601,014 | 289 bp | NM_021804 |
| dACE2, Ex1c | GTAATTCCCAGGTTGCAGGCTT<br>GTGAGAGCCTTAGGTTGGATTC<br>CCTAGCTTGAAAAGGAGATCGT<br>TTTACAAGTGCTTCATTGAGGA<br>GAGCTCTGAGGCAGAGGGGAA<br>TGAGGGAAGCAGGCTGGGACA<br>AAGGAGGGAG | chrX:15,580,281-15,580,420 | 140 bp | MT505392 |

### Supplementary Figures

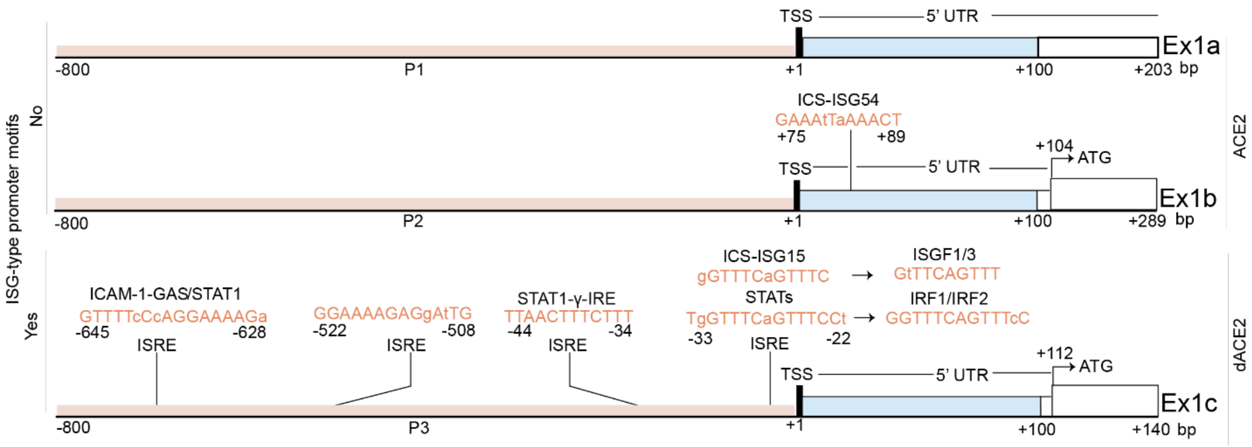


**Figure S1. Analysis of promoter regulatory elements relevant for IFN signaling**

Promoters of *ACE2* (P1 and P2) and *dACE2* (P3) were analyzed for binding motifs of transcription factors relevant for IFN signaling. Promoters were defined within the -800 bp/+100 bp window from the corresponding transcription start sites (TSS).

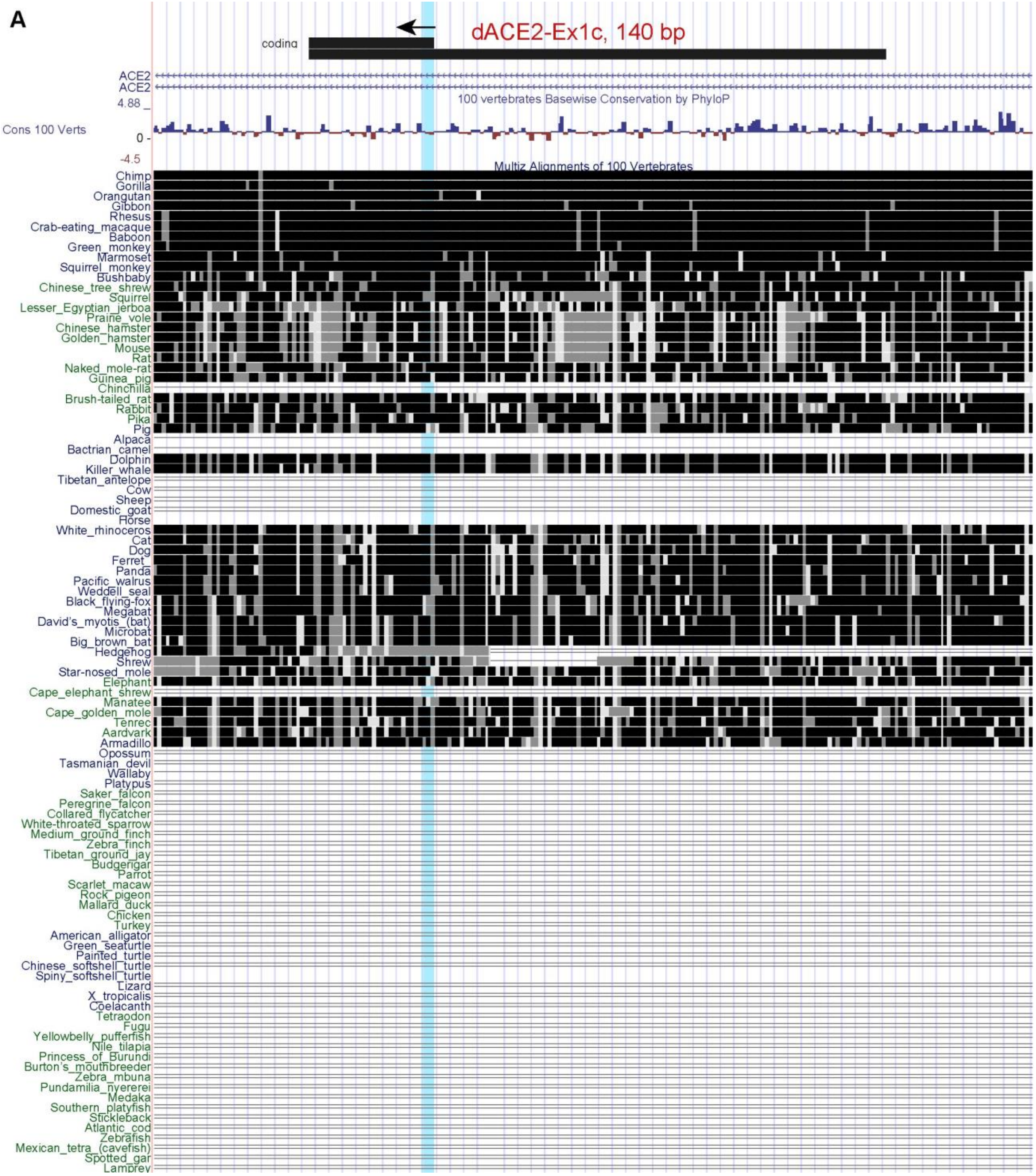
A
coding
dACE2-Ex1c, 140 bp
ACE2
ACE2
4.88 _
100 vertebrates Basewise Conservation by PhyloP
Cons 100 Verts
0 -
-4.5
Multiz Alignments of 100 Vertebrates
Chimp
Gorilla
Orangutan
Gibbon
Rhesus
Crab-eating_macaque
Baboon
Green_monkey
Marmoset
Squirrel_monkey
Bushbaby
Chinese_tree_shrew
Squirrel
Lesser_Egyptian_jerboa
Prairie_vole
Chinese_hamster
Golden_hamster
Mouse
Rat
Naked_mole-rat
Guinea_pig
Chinchilla
Brush-tailed_rat
Rabbit
Pika
Pig
Alpaca
Bactrian_camel
Dolphin
Killer_whale
Tibetan_antelope
Cow
Sheep
Domestic_goat
Horse
White_rhinoceros
Cat
Dog
Ferret_
Panda
Pacific_walrus
Weddell_seal
Black_flying-fox
Megabat
David's_myotis_(bat)
Microbat
Big_brown_bat
Hedgehog
Shrew
Star-nosed_mole
Elephant
Cape_elephant_shrew
Manatee
Cape_golden_mole
Tenrec
Aardvark
Armadillo
Opossum
Tasmanian_devil
Wallaby
Platypus
Saker_falcon
Peregrine_falcon
Collared_flycatcher
White-throated_sparrow
Medium_ground_finch
Zebra_finch
Tibetan_ground_jay
Budgerigar
Parrot
Scarlet_macaw
Rock_pigeon
Mallard_duck
Chicken
Turkey
American_alligator
Green_seaturtle
Painted_turtle
Chinese_softshell_turtle
Spiny_softshell_turtle
Lizard
X_tropicalis
Coelacanth
Tetraodon
Fugu
Yellowbelly_pufferfish
Nile_tilapia
Princess_of_Burundi
Burton's_mouthbreeder
Zebra_mbuna
Pundamilia_nyererei
Medaka
Southern_platyfish
Stickleback
Atlantic_cod
Zebrafish
Mexican_tetra_(cavefish)
Spotted_gar
Lamprey

B

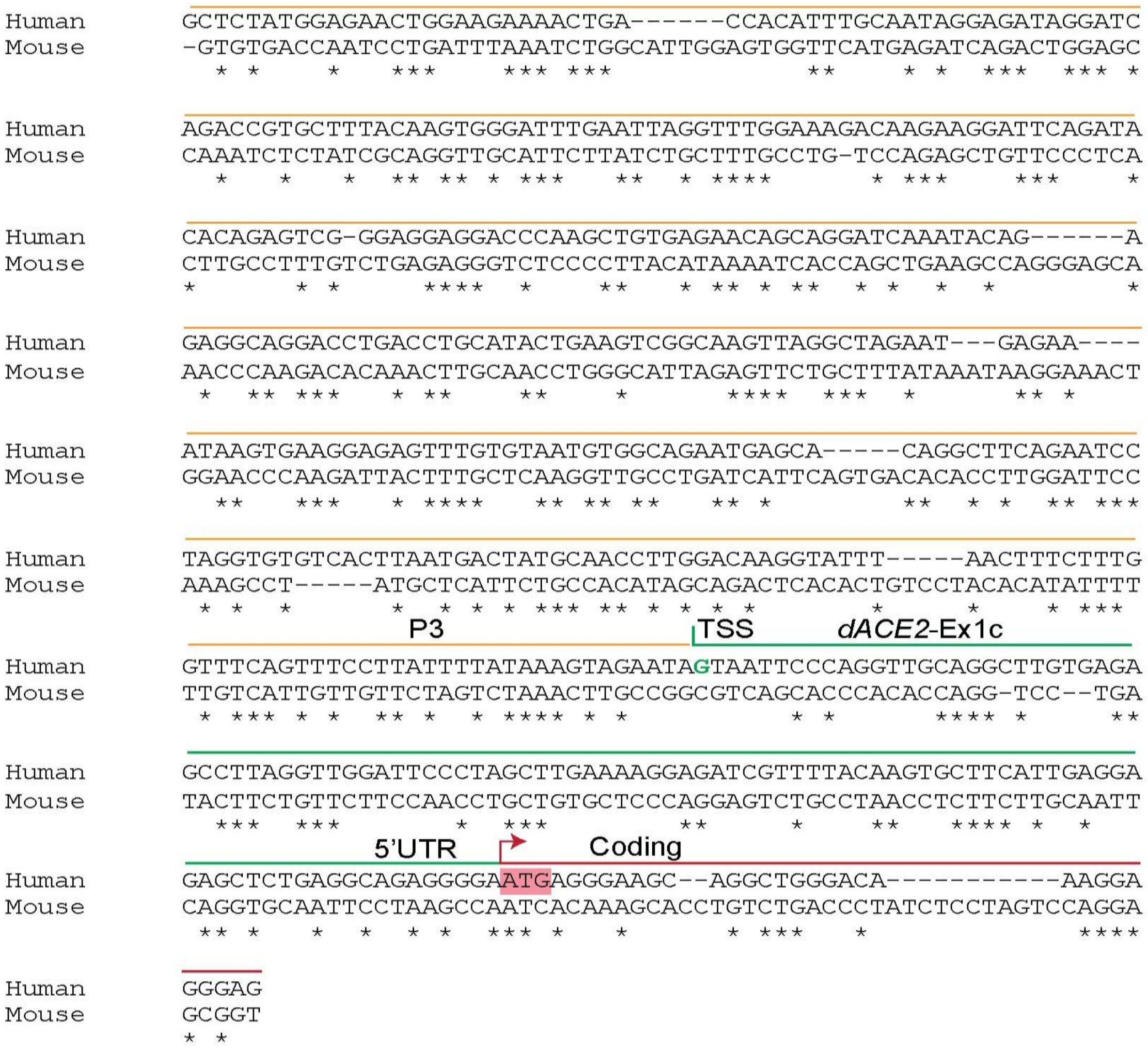


**Figure S2. Conservation of the *dACE2*-Ex1c sequences.**

**A**). Conservation of the 140 bp sequence of *dACE2*-Ex1c (human chrX:15,580,281-15,580,420, GRCh38/hg38) was analyzed by BLAT in 100 vertebrate species in the UCSC genome browser (www.genome.ucsc.edu). The sequence is highly conserved in primates but is less conserved or absent in non-primates, precluding *dACE2* transcript initiation or translation into an ACE2-type protein. The long bar indicates the entire Ex1c (140 bp) and the short bar indicates the protein-coding part of this exon (30 bp), starting from the ATG codon indicated by an arrow and highlight; the gene direction is from right to left. **B**). Comparison between human and mouse sequences; *- conserved bases; transcription start site (TSS) and translation start site (ATG) are indicated based on the human sequence. Human and mouse sequences share 43.7% identity within 500 bp (includes Ex1c, 5’UTR and promoter). Sequences were downloaded from UCSC genome browser and aligned using Clustal Omega.

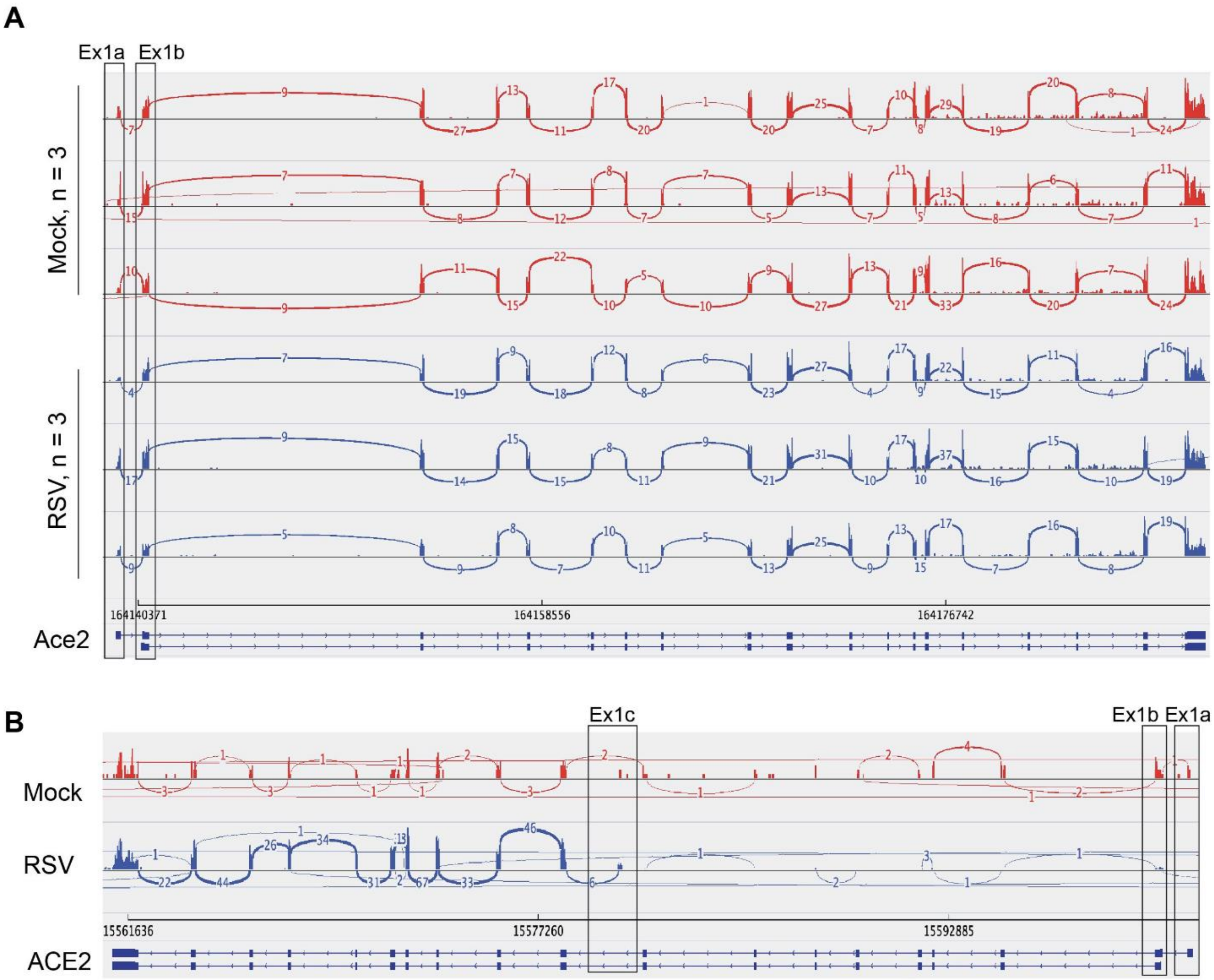


**Figure S3.** ***ACE2*** **expression patterns in mouse and human lung cells infected with the respiratory syncytial virus (RSV).**

**A**) Sashimi plots of the *Ace2* region in a lung RNA-seq dataset from mice mock/RSV- infected (in triplicates). *Ace2*-Ex1a and Ex1b show similar expression patterns in all samples. The expression of *dACE2*-Ex1c is not observed, consistent with the absence of the corresponding genomic sequence in mice (**Figure 1D**, **Figure S1A, B**). **B**) Sashimi plots of the *ACE2* region in H292, a human lung mucoepidermoid pulmonary carcinoma cell line, show that expression of *ACE2* from Ex1a and Ex1b and *dACE2* from Ex1c is very low at baseline. Only *dACE2* expression is induced by RSV infection. Note: The mouse and human *ACE2* genes are shown in opposite orientations, as presented in the Integrative Genomic Viewer (IGV). Dataset: PRJNA588982.

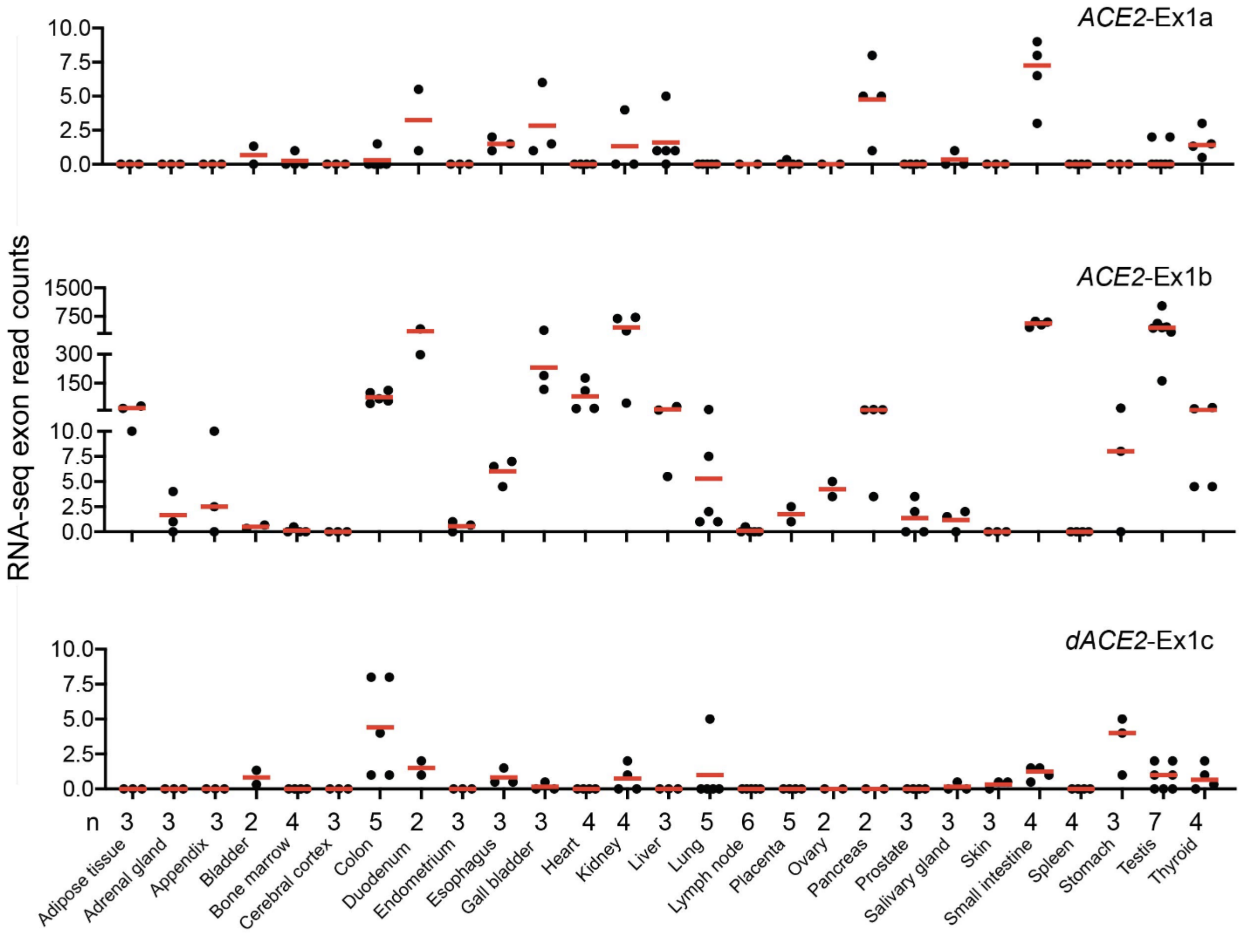


**Figure S4. Expression of *ACE2* and *dACE2* in normal human tissues.** RNA-seq read counts for *ACE2*-Ex1a and Ex1b and *dACE2*-Ex1c in 27 human tissues. Dataset: PRJEB4337, n = 95

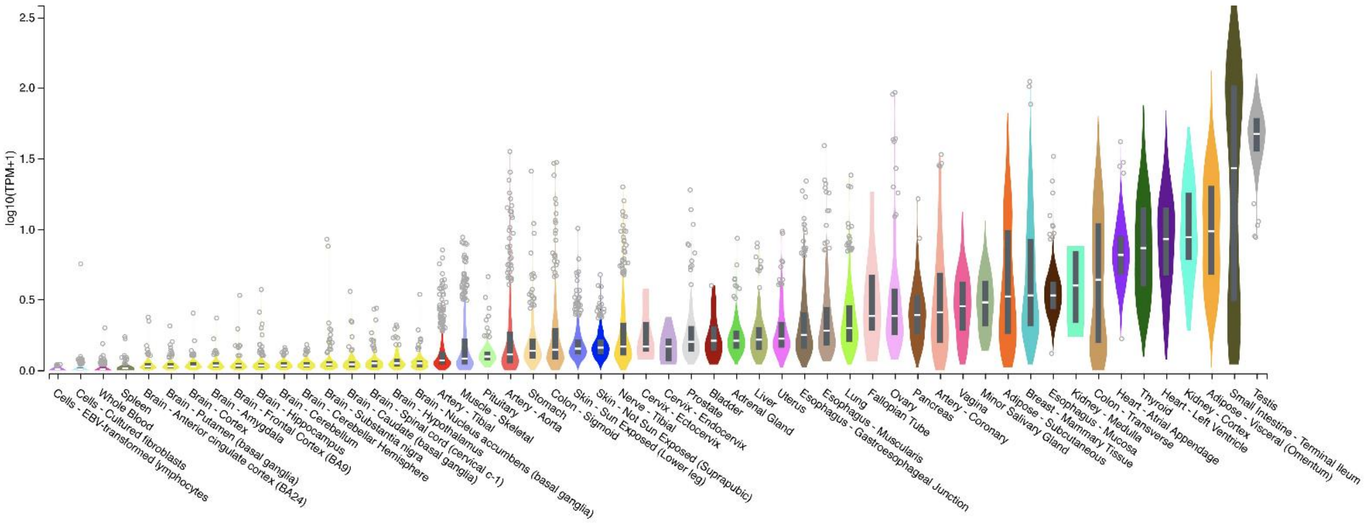


**Figure S5.** ***ACE2*** **expression in the Genotype-Tissue Expression (GTEx) project.**

Gene-based *ACE2* expression (combines *ACE2* and *dACE2* isoforms) in 17,382 normal human tissue samples of 54 tissue types in GTEx https://www.gtexportal.org/home/gene/ACE2.

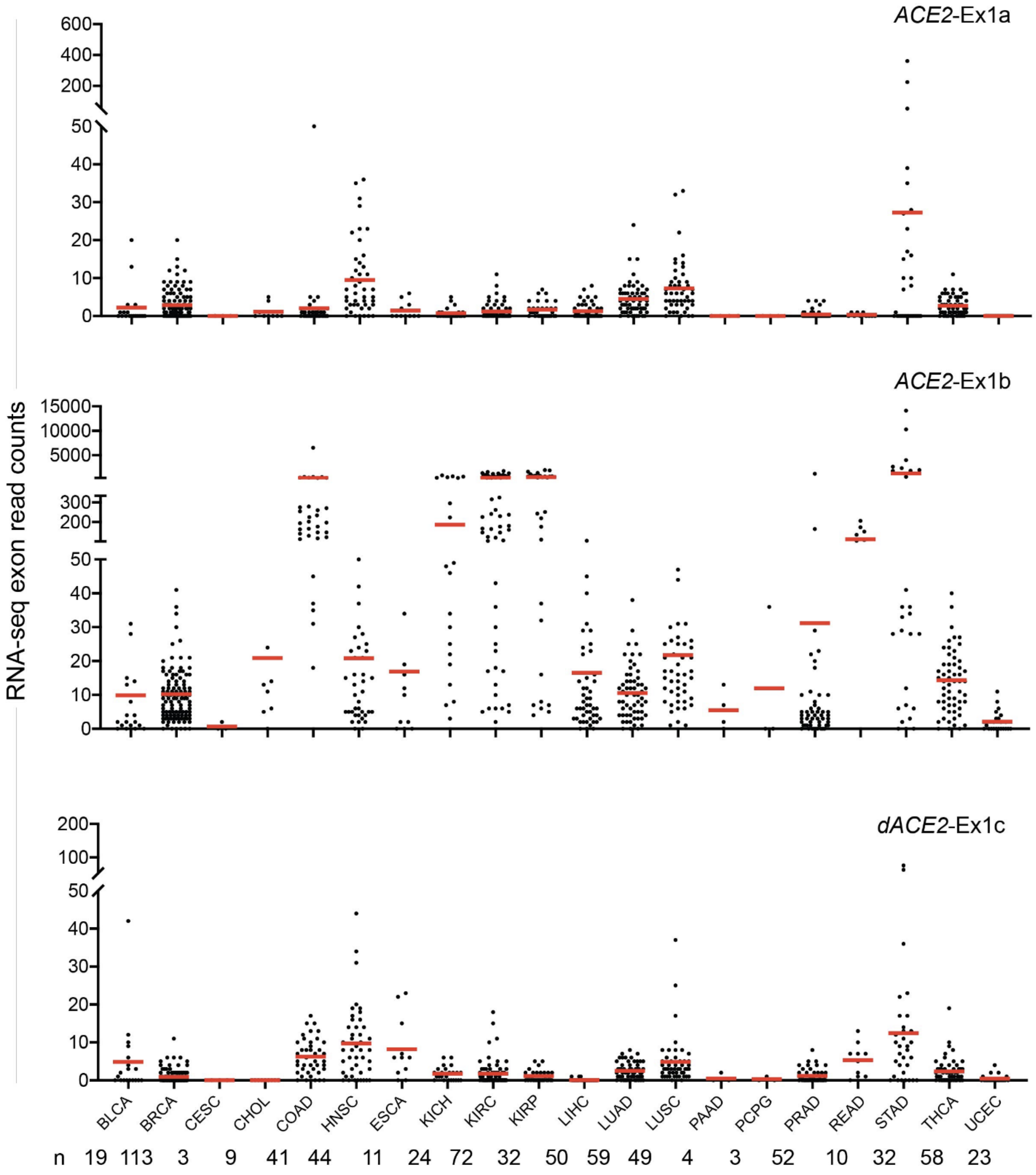


**Figure S6. Expression of *ACE2* and *dACE2* in tumor-adjacent normal tissues in TCGA.** Based on RNA-seq read counts, *ACE2*-Ex1b is detectable in multiple samples of several tissue types. *dACE2*-Ex1c expression is more restricted and most common in normal tissue adjacent to tumors of head and neck (HNSC), stomach (STAD), lung squamous carcinoma (LUSC), colon (COAD), and esophagus (ESCA).

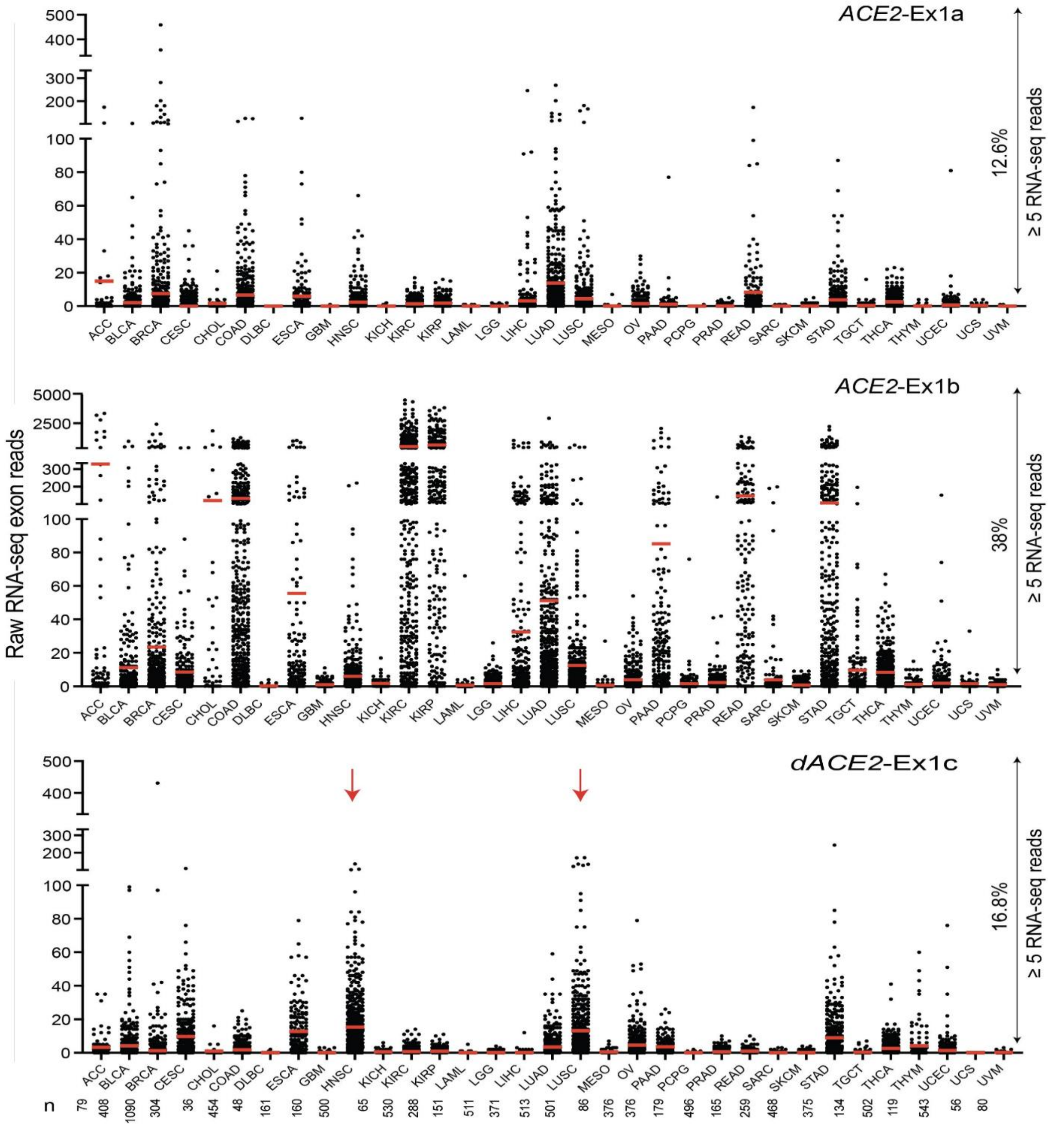


**Figure S7. Expression of *ACE2* and *dACE2* across 10,185 tumors of 33 types in TCGA.** Based on RNA-seq read counts, *ACE2*-Ex1b is most expressed in kidney tumors - kidney renal clear cell carcinoma (KIRC) and kidney renal papillary cell carcinoma (KIRP). Most samples expressing *dACE2*-Ex1c are squamous tumors of head and neck (HNSC) and the lungs (LUSC). Based on ≥5 reads/sample threshold, *ACE2*-Ex1a is expressed in 12.6%, *ACE2*-Ex1b – in 38.0% and *dACE2*-Ex1c - in 16.8% of all tumors.

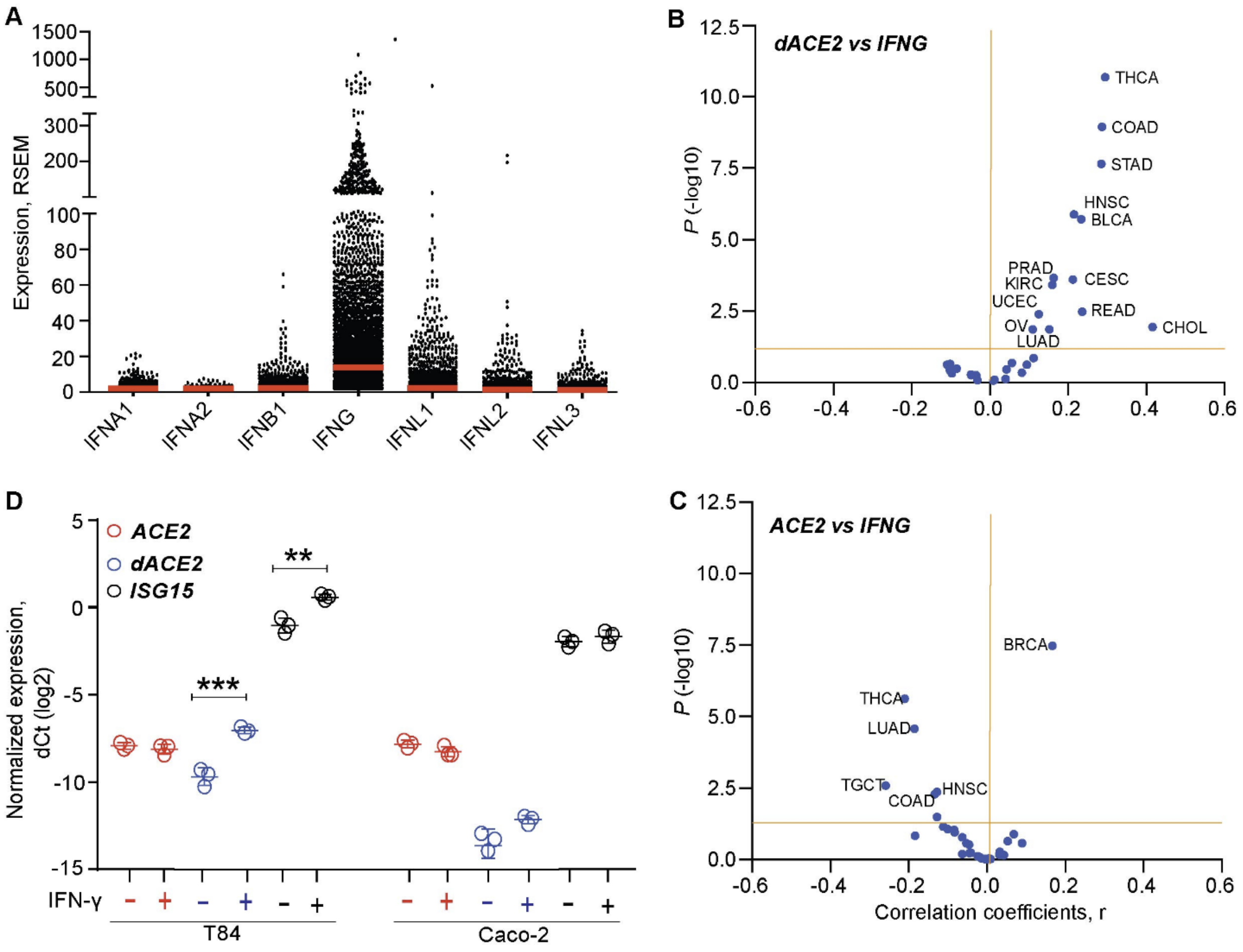


**Figure S8. Analysis of *dACE2* and *ACE2* expression in relation to *IFNG* expression in TCGA tumors and *in vitro* IFN-ɣ treatment.**

**A)** Expression levels of all *IFN* genes annotated in TCGA tumors (n = 10,185) were acquired from cBioPortal (https://www.cbioportal.org/); expression of *IFNL4* was not available. At RSEM ≥ 1, only expression of *IFNG* is common (61% samples), with mean expression RSEM=19.8 compared to other IFN genes (mean expression RSEM ≤ 1.3). **B**, **C)** Pearson correlation coefficients (r) for *dACE2* and *ACE2* vs. *IFNG* expression across tumors. *dACE2* showed significant positive correlations (r ≥ 0.2) with *IFNG* in 8 tumor types, while *ACE2* showed mainly negative correlations and only one positive correlation in breast cancer (r = 0.15). Expression values for *dACE2* and *ACE2* were based on log2 normalized exon read counts (Ex1b and Ex1c) and for *IFNG* - on RSEM values. **D)** Treatment of cell lines with IFN-ɣ (2ng/ml, 48 hrs) induced expression of *dACE2* but not *ACE2* in T84 cells.

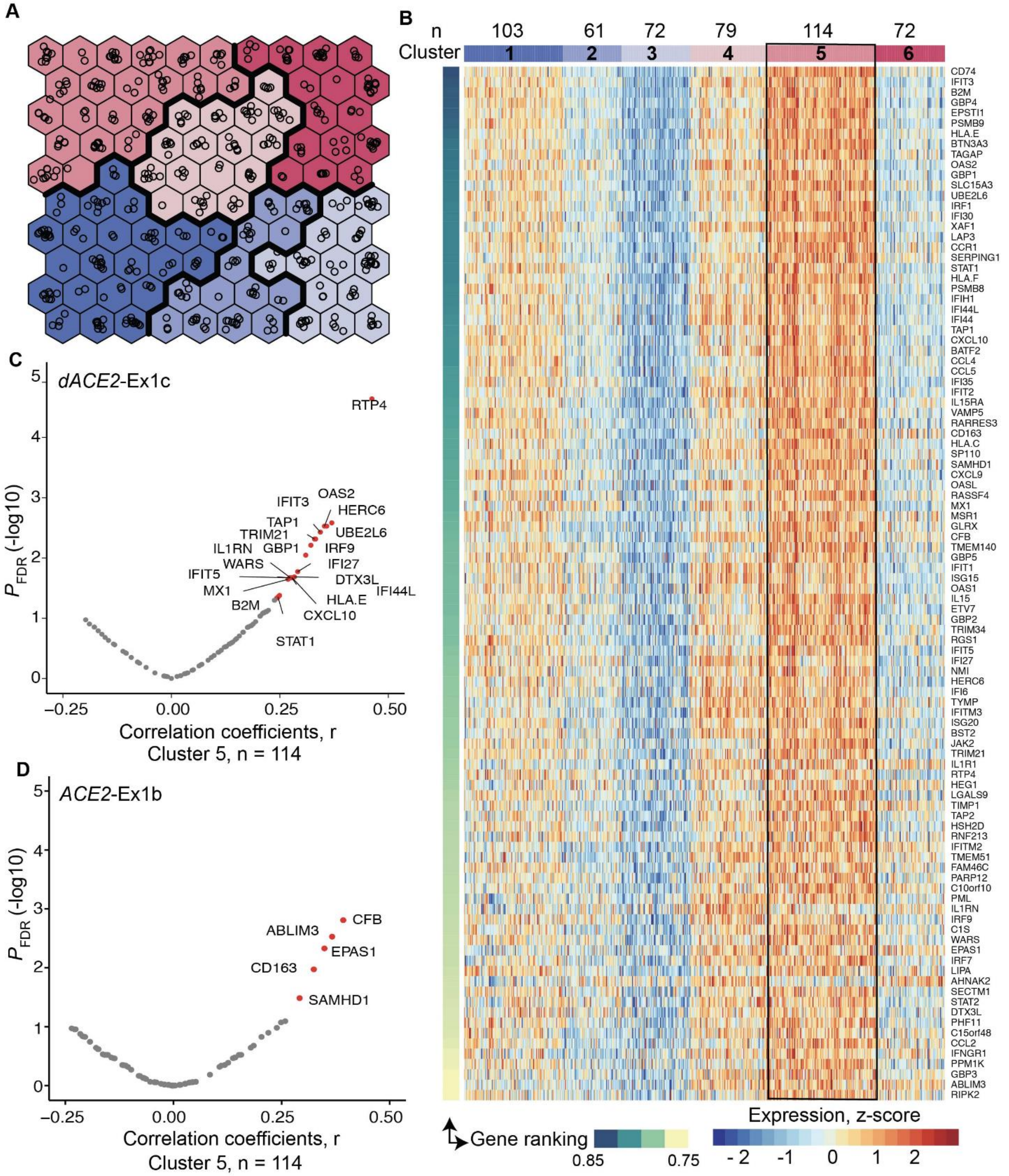


**Figure S9. Unsupervised self-organizing map (SOM) analysis in TCGA-LUSC tumors**

**A)** Construction of the unsupervised SOM of TCGA-LUSC tumors (n=501) based on Z-scores calculated for each of the 270 curated ISGs. Each hexagon includes a mean of 5 (range 1-14) tumors with similar ISG expression profiles. Colors denote clusters (1-6) of tumors with similar ISG expression profiles. **B)** Heatmap of the 6 SOM-defined clusters plotting the expression of top 100 ISGs selected by ranking of the initial set of 270 ISGs based on their contribution to these clusters. Cluster 5 includes 114 tumors with the highest ISG expression, whereas cluster 3 includes 72 tumors with the lowest ISG expression. **C)** Volcano plots showing FDR-adjusted p-values and Pearson correlation coefficients (r) for expression of *dACE2* and *ACE2* in relation to expression of the top 100 ISGs within cluster 5. In total, *dACE2* was significantly (FDR p-value $< 0.05$) correlated with expression of 20 ISGs and *ACE2* - with 5 ISGs.

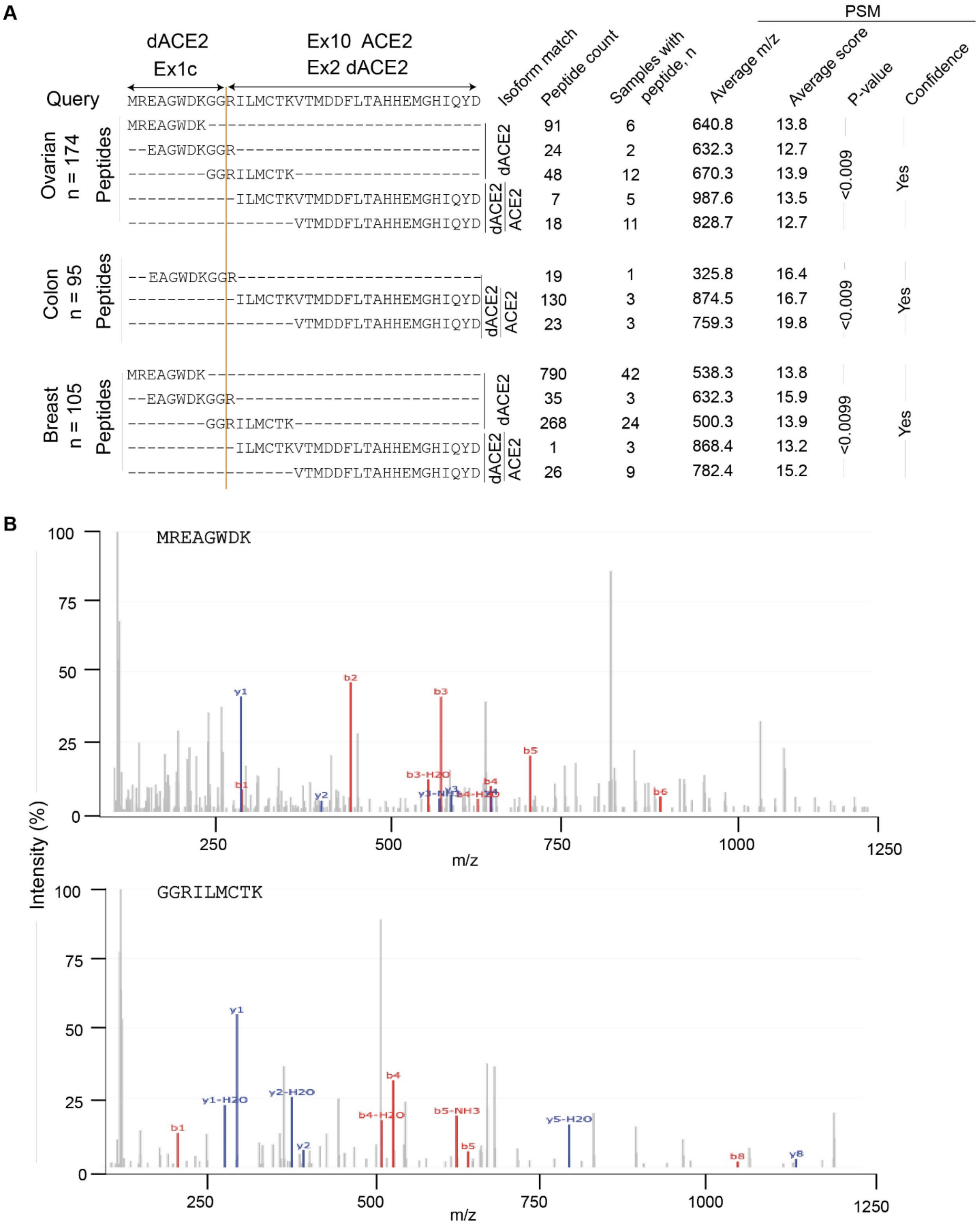


**Figure S10. Peptides encoded by d*ACE2*-Ex1c are detected by protein sequencing in tumors.**

**A)** Results of peptide query in PepQuery2 proteomics database of mass-spec data in 174 ovarian, 95 colon, and 105 breast tumors in TCGA (Wen et al., 2019). Three peptides – MREAGWDK, EAGWDKGGR, and GGRILMCTK uniquely correspond to 10 aa encoded by *dACE2*-Ex1c. The latter peptide results from the splicing of *dAC2*-Ex1c with its downstream exon. The total number of identified peptides, the number of samples with specific peptides, and corresponding parameters for a peptide-spectrum match (PSM) are shown in table format. **B)** Representative spectra of two peptides matching with the protein encoded by *dACE2*-Ex1c. M/z refers to the mass by charge ratio. The b-series and y-series ions showed the correct mapping of residues in the query aa sequence.
